# Fate and state transitions during human blood vessel organoid development

**DOI:** 10.1101/2022.03.23.485329

**Authors:** Marina T. Nikolova, Zhisong He, Reiner A. Wimmer, Makiko Seimiya, Jonas M. Nikoloff, Josef M. Penninger, J. Gray Camp, Barbara Treutlein

## Abstract

Blood vessel organoids (BVOs) derived from human pluripotent stem cells have emerged as a novel system to understand human vascular development, model disorders, and develop regenerative therapies. However, it is unclear which molecular states constitute BVOs and how cells differentiate and self-organize within BVOs *in vitro* and after transplantation. Here we reconstruct BVO development over a time course using single-cell transcriptomics. We observe progenitor states that bifurcate into endothelial and mural fates, and find that BVOs do not acquire definitive arterio-venous endothelial identities *in vitro*. Chromatin accessibility profiling identifies gene regulatory network (GRN) features associated with endothelial and mural fate decisions, and transcriptome-coupled lineage recording reveals multipotent progenitor states within BVOs. We perform single-cell genetic perturbations within mosaic BVOs to dissect the impact of transcription factor (TF) and receptor depletion on cell differentiation, and highlight multiple TFs including MECOM and ETV2 as strong-effect regulators of human BVO development. We show that manipulation of VEGF and Notch signaling pathways alters BVO morphogenesis and endothelial GRNs, and induces arteriovenous-like state differentiation. We analyze matured BVOs after transplantation using scRNA-seq, and observe matured endothelium with clear arteriovenous specification. We also observe off-target cell fates with bone and adipocyte features, suggesting multipotent states reside within the BVOs *in vitro* that expand and diversify in less restrictive conditions. Finally, we map vascular disease associated genes to BVO cell states to highlight the potential of BVOs for disease modeling. Altogether, our data and analyses provide the first comprehensive cell state atlas of BVO development and illuminate both the power and limitation of BVOs for translational research.

## Introduction

The vasculature is the first organ to form and is essential to the proper development and function of all other tissues in the human body (Gilbert, 2010). Vascular dysfunction has a major role in the initialization and progression of multiple disabling and life-threatening conditions. In order to create accurate models of human disease pathophysiology and engineer tissue therapies, it is key to understand the complex processes of blood vessel formation and recapitulate their biological functions using human cells. During development, endothelial cell (EC) progenitors, termed angioblasts, arise from the mesoderm and aggregate to form a primitive vascular plexus (Ferguson et al., 2005; Okuda and Hogan, 2020). Subsequently, during angiogenesis, these primitive blood vessels guided by tip cells (specialized ECs at the extremity of blood vessels guiding migration) degrade surrounding basement membranes, sprout, form tubules and connect into mature circulatory networks (Okuda and Hogan, 2020). Endothelial cells synthesize a new basement membrane and recruit pericytes and smooth muscle cells (SMCs) (associated with small-and large-diameter vessels, respectively; collectively known as mural cells) and perivascular fibroblasts. Mural cells have organ-specific origins (Creazzo et al., 1998; Korn et al., 2002; Que et al., 2008; Wasteson et al., 2008), and are critical for blood vessel stability, structure and barrier formation (Sweeney and Foldes, 2018). An instructive role in the development and specification of the vasculature is played by manifold signaling pathways, with Notch, VEGF, TGF-β, Angiopoietin-Tie and BMP being some of the most prominent (Schmidt and Liebner, 2015). The VEGF pathway has proven indispensable for early vasculature development and mice lacking the receptor VEGFR2 (encoded by KDR in humans) or the ligand VEGF-A die in utero as a result of complete failure to form blood vessels (Carmeliet et al., 1996; Ferrara et al., 1996; Shalaby et al., 1995). Furthermore, the crosstalk between VEGF and Notch signaling regulates sprouting angiogenesis and differentiation into arteries, veins and capillaries (Fang and Hirschi, 2019; Mack and Luisa Iruela-Arispe, 2018). Thus, many clinically approved anti-angiogenic agents, such as those in chemotherapy and retinal vascular pathologies, selectively inhibit the activity of VEGF-A or Notch alone or in combination with other pathways (Akil et al., 2021; Christopoulos et al., 2021; Ghanchi et al., 2022). The exact transcriptional mechanisms through which gene expression downstream of these signaling pathways is regulated remain to be fully elucidated.

A multitude of transcription factors (TFs) have emerged as critical regulators of endothelial and hematopoietic cell development. ETV2, a member of the ETS TF family, is transiently expressed in mesodermal progenitors giving rise to cardiovascular lineages and has been identified as indispensable for EC fate (Kataoka et al., 2011; Lee et al., 2008). Groups have reported direct reprogramming of human fibroblasts into ECs by overexpression of ETV2 alone (Lee et al., 2017; Morita et al., 2015). Other ETS members such as FLI1 and ERG, as well as GATA2, SOX7, SOX18, NR2F2 and others have been recognized as major regulators of EC development, function, angiogenic growth and arteriovenous specification (Craig and Sumanas, 2016; Yao et al., 2019; You et al., 2005). The precise definition and ontogeny of mural and perivascular cells, however, have been widely debated due to the differences in their embryonic origin between tissues and within the same tissue (Dias Moura Prazeres et al., 2017). Therefore much less is known about the transcriptional regulation of mural cell development, but recent reports have indicated the importance of EBF1, NKX3.2 for pericyte development in different organs (Harari et al., 2019; Pagani et al., 2021). Most knowledge about the signaling pathways and transcription factors that regulate vascular development has been derived from chick, zebrafish and mouse models. New human model systems are needed to understand and explore the regulatory networks underlying human vasculature development.

Recent breakthroughs in stem cell culture have led to the development of protocols for human pluripotent stem cell (hPSC)-derived blood vessel models, which have potential for gaining insight into human angiogenesis, disease modeling, and regenerative therapy. Examples include mono-and co-culture of hPSC-derived endothelial and perivascular cells, which are capable of forming vessel-like structures upon embedding in a matrix (Kusuma et al., 2013; Orlova et al., 2014). Recently, a protocol was developed to generate human blood vessel organoids (hBVOs) (Wimmer et al., 2019a), which are self-organizing three-dimensional structures that recapitulate certain steps of human vasculature development *in vitro*. After mesoderm induction, vessels sprout, networks form, and subsequently develop approximate aspects of human vasculature development *in vivo* (Wimmer et al., 2019a). The hBVO system offers great promise for modeling human disease (Monteil et al., 2020; Wimmer et al., 2019b), and as a tool for therapy development and more accurate or complex tissue engineering (Ahn et al., 2021; Dailamy et al., 2021). However, we lack a detailed cell state atlas across hBVO development, and we do not yet understand the power and limitations of this system in order to harness its full potential.

Here, we use single-cell RNA sequencing (scRNA-seq), single-cell chromatin accessibility sequencing (scATAC-seq), and whole-mount immunohistochemistry to characterize human blood vessel organoid development. We explore endothelial and mural cell fate bifurcation utilising single-cell transcriptome coupled-lineage recording and multiplexed *in organoid* CRISPR perturbation and identify effects on cell fate decisions during organoid differentiation. We show that upon transplantation into immunocompromised mice, endothelial cells mature and acquire arterial-or venous-like phenotypes. We modulate VEGF and Notch pathways *in vitro* to explore hBVO response to chemical perturbation and identify gene expression alterations involved in arteriovenous specification and vascular sprouting. Finally, we present a disease-associated gene expression catalog of *in vitro* and transplanted human vascular organoids, and published primary endothelial cells across organ systems. Altogether, we present a comprehensive single-cell genomic atlas of developing hBVOs and their perturbed states, which can serve as an extensive resource for studying human vascular development and disease and lay the groundwork for future BVO engineering efforts.

## Results

### Single-cell transcriptome analysis of human blood vessel organoid development

We generated hBVOs following an adapted version of the original protocol (Wimmer et al., 2019a), where mesoderm aggregates were grown in commercially available microwell plates prior to embedding in a Matrigel/Collagen-I matrix (Fig. 1A). Immunohistochemical analysis of developing hBVOs (Fig.1B; Fig. S1) revealed PDGFR-β^+^ mesodermal cells at day 3 of differentiation and the emergence of CD31^+^ endothelial cells in the core of mesoderm aggregates at day 4, both of which remained prevalent in the later stages of organoid development (Fig. 1B; Fig. S1). At day 6, one day after matrix embedding, PDGFR-β^+^ and CD31^+^ cells sprouted from the aggregates and formed vascular networks (Fig. 1B; Fig. S1D). In the network periphery, where active angiogenesis occurs (angiogenic front), endothelial cells marked by the VEGFR co-receptor NRP2 (Alghamdi et al., 2020) and enriched in APLNR (Helker et al., 2020) appeared to lead the vessel formation and exhibited tip cell morphology with extended filopodia (Fig. 1C; Fig. S1H). At day 10, we detected the formation of vessel lumens, lined by endothelial cells and surrounded by mural cells (Fig. 1D; Fig. S1I). Upon isolation from the matrix, networks were transferred to an ultra-low attachment plate, where they self-assembled into blood vessel organoids consisting of endothelial and mural cells ensheathed by a basement membrane marked by Collagen IV (Fig. 1E; Fig. S1F, G, J).

**Figure 1.**
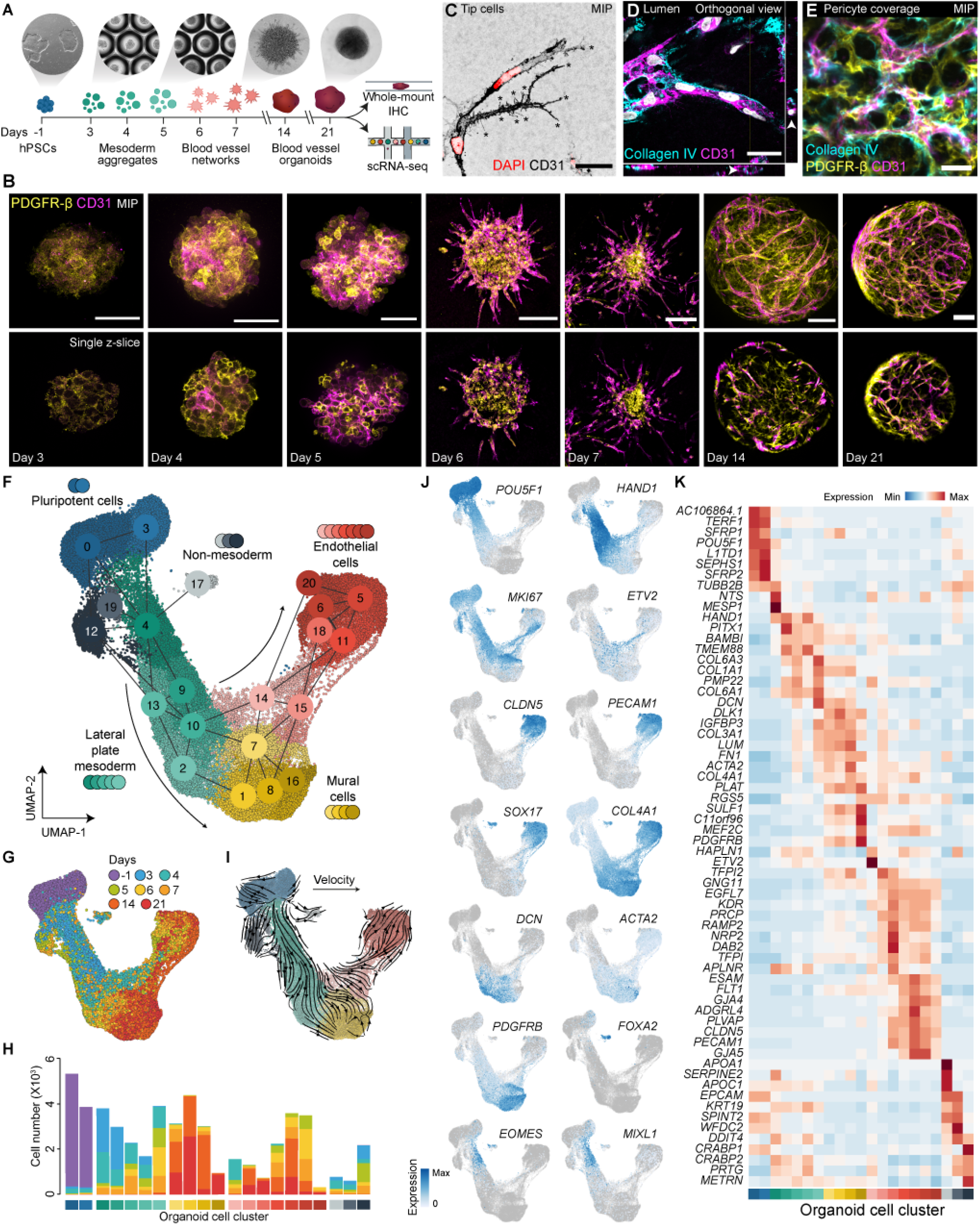
Reconstructing human blood vessel organoid differentiation from pluripotency using single-cell transcriptomics and immunohistochemistry. (A) Schematic of the hBVO protocol. scRNA-seq, single-cell RNA-sequencing; IHC, immunohistochemistry. (B) IHC of hBVOs throughout differentiation. PDGFR-β (yellow) marks mural cells; CD31 (magenta) marks endothelial cells. Scale bars: 50 µm (day 3, 4, 5), 100 µm (day 6, 14), 150 µm (day7), 200 µm (day 21). MIP: maximum intensity projection. (C) IHC at day 7 detecting CD31 (black) and DAPI (nuclei; red) endothelial tip cells. Asteriks, filopodia. Scale bar: 30 µm. (D) Orthogonal view of endothelial cells marked by CD31 (magenta), embedded in a basement membrane (Collagen IV, cyan) forming a lumen (arrowheads). Scale bar: 30 µm. DAPI, nuclei, white. (E) Endothelial cell network (CD31, magenta) covered by mural cells (PDGFR-β), embedded in a basement membrane (Collagen IV, cyan). Scale bar: 20 µm. (F, G) UMAP embedding of scRNA-seq data with cells colored, labeled by cell type cluster (F) or by time point (G). (H) Number of cells (y-axis) in each cluster (x-axis) at each time point (bar color). (I) Streamline plot of a UMAP-based embedding colored by cluster and labeled with vector fields from RNA velocity (scVelo). (J) Feature plots of cell type cluster marker genes across the trajectory. (K) Heatmap showing the scaled average marker gene expression (rows) across clusters (columns). See also figures S1 to S3.

Next, we used single-cell RNA-sequencing (scRNA-seq; 10X Genomics) to characterize the cell composition across hBVO differentiation using human embryonic stem cells (hESCs; H9) and induced pluripotent stem cells (hiPSCs; NC8) (Fig. S2A). We sequenced RNA from 52,680 cells at eight time points (stem cells -1, 3, 4, 5, 6, 7, 14 and 21 days of differentiation, pooling 20-30 aggregates or 15-20 organoids per time point) (Table S1). At each time point, we combined and dissociated tissues from both cell lines together, and subsequently demultiplexed the sequencing data using single nucleotide polymorphisms (SNPs) differing between the two individuals (demuxlet (Kang et al., 2018)). After quality control, we integrated the datasets using cluster similarity spectrum (CSS (He et al., 2020)) and visualized cell transcriptomes in a Uniform Manifold Approximation and Projection (UMAP (McInnes et al., 2018)) embedding (Fig. 1F; Fig. S2B). Heterogeneity analysis resulted in 21 molecularly distinct clusters, which we annotated based on cluster-enriched genes identified by differential expression analysis (Table S2).

Based on markers and similarity to cells from a published developing human multi-organ cell atlas (Yu et al., 2021), we annotated clusters as “pluripotent cells” (c0, c3), “lateral plate mesoderm” (c4, c9, c13, c10, c2), “mural cells” (c7, c1, c8, c6), “endothelial cells” (ECs) (c14, c15, c18, c11, c5, c6, c20) and “non-mesoderm cells” (c17, c19, c12) (Fig. 1F-J; Fig. S2C). We observed a relatively continuous trajectory from pluripotency through a bifurcation into endothelial and mural cell lineages. In correlation with tissue age (Fig. 1G), stem cells (POU5F1^+^; c0, c3) progressed through a primitive streak/early mesoderm-like stage, marked by the expression of EOMES, MIXL1, NTS and MESP2, at day 3 (c4; Fig. 1J,K; Fig. S2E), into lateral plate mesoderm cells expressing HAND1, PITX1 and BAMBI at days 3-7 (c9, c13, c10, c2) (full gene names in Table S3). We note three small non-mesoderm populations (c17, c19, c12), which appeared to not progress further in the hBVO differentiation and were not apparent at and after day 14.

Expression of the endothelial cell development master regulator ETV2 peaked in a subpopulation of cells at day 4 (c14) and decreased in expression in later stages of differentiation (Fig. 1J, K). Based on previous studies, RNA velocity analysis using spliced and unspliced transcripts, and inspection of marker genes, we hypothesized that these ETV2^+^ cells marked an intermediate stage during endothelial cell maturation and therefore called them early endothelial progenitor cells (EPCs). Further EC differentiation was marked by the upregulation of KDR, NRP2, CLDN5 and PECAM1 (c18, c11, c5, c6, c20) at days 5-21 (Fig. 1J, K). We did not observe any further increase in EC marker gene expression between day 14 and 21. However, we could identify a strong induction of venous EC markers at day 21 (c18), while arterial-like EC populations largely consisted of cells from hBVOs at days 6, 7 and 14 (Fig. S2D).

Other mesoderm cell clusters (c7, c1, c8, c16) showed expression of DCN, DLK1, LUM, ACTA2 and PDGFRB, known to mark fibroblasts, SMCs and pericytes and we collectively call these cells mural cells. The majority of mural cells consisted of cells from day 14 and 21 hBVOs and were enriched in the heparan sulfate endosulfatase SULF1, which has been associated with inhibition of EC angiogenesis (Narita et al., 2006). A gene expressed by both mural and ECs was COL4A1, an essential component of the vascular basement membrane (Fig. 1J, K). Immunohistochemical analysis confirmed the deposition of COL4A1+ basement membrane early in hBVO development (Fig. S1A-G).

Interestingly, we observed two differentiation trajectories towards endothelial cells that separated by time point. In addition to the differentiation of lateral plate mesoderm progenitor cells into ECs through EPC cluster 14 at days 4-7, there was a second branch of EPCs (late EPCs; c15) mostly present in organoids at day 14 and 21 and forming a trajectory from differentiated mural cells towards ECs (Fig. 1H, J). This cluster did not express ETV2 and instead was characterized by the expression of TFPI2, as well as both mural and EC markers such as LUM and NRP2, respectively. Reconstruction of an EC differentiation pseudotime supported the hypothesis that two distinct EC progenitor populations develop into ECs (Fig. S2E-H). Altogether, these data provide a detailed assessment of cell heterogeneity and differentiation trajectories in developing hBVOs.

### Lineage recording highlights the potential of hBVO mural cells to differentiate into endothelial lineages

To further explore the heterogeneity within EC progenitors (c14 and c15; Fig. S3A-D) and their differentiation potential, we utilized iTracer (He et al., 2022) an inducible lineage recording system based on cell reporter barcoding, CRISPR/Cas9 scarring and single-cell transcriptomics to resolve lineage relationships. The iTracer construct contains a poly-adenylated and barcoded (11 random bases) red fluorescent protein (RFP) driven by the RPBSA promoter as well as a hU6 promoter-driven gRNA targeting a region within the 3’ portion of the RFP coding sequence (Fig. S3E). We transfected a doxycycline-inducible Cas9 (iCas9) hiPSC line (González et al., 2014; Riesenberg and Maricic, 2018) with the construct, so that each transfected cell contained at least one barcode, then expanded and isolated RFP+ cells using fluorescence activated cell sorting (FACS) (Fig. S3F). Next, we generated hBVOs and induced transient Cas9 expression (Fig. S3G) by doxycycline treatment at day 7, a time point at which the late EPCs (c15) are the predominant endothelial progenitor population. This caused the formation of Cas9-gRNA complexes creating insertions and/or deletions (scars) at the targeted location in the RFP region. The scars together with the unique barcodes are inherited within cell lineages and can be detected by sequencing, serving as a powerful dual channel lineage recorder.

At day 14 of hBVO differentiation, we performed scRNA-seq of 4,948 cells with targeted amplicon enrichment of barcode and scar regions. Based on marker gene analysis, we annotated these cells as either mural or endothelial cells (Fig. S3H). We detected barcode and RFP sequences in 1,924 cells, 898 of which possessed one or more scars (Table S1). We found 20 cell groups, formed by at least ten cells, sharing the same barcode composition and hence deriving from the same pluripotent stem cell (barcode families; Fig. S3I). Ten of those barcode families contained at least three cells sharing the same scar composition and hence originating from the same cell in the BVO at day 7, the time point of scarring (scar families; Fig. S3J). All scar families were either composed of only mural cells, or mural and endothelial cells, suggesting that indeed cells in hBVOs at day 7 still have the potential to differentiate into both mural and endothelial cells. These experiments provide a set of tools to explore endothelial and mural cell lineage relationships, and confirm the potency of cells within the early hBVOs.

### Paired single-cell RNA and chromatin accessibility profiling reveals candidate regulators of endothelial and mural cell fate maintenance

To identify regulatory mechanisms controlling endothelial and mural cell fate decisions, we performed paired scRNA-seq and scATAC-seq from the same nuclei (16,648 nuclei, scMultiome, 10X Genomics) of two cell lines (H9, a hESC and NC8, a hiPSC line) at day 7 of hBVO differentiation (Fig. 2A; Table S1). We applied CSS to integrate the two cell lines and generated a UMAP embedding of the transcriptome and the accessible chromatin landscapes (Fig. S4A). We then computed a joint weighted nearest neighbor (WNN) graph, representing both the gene expression and DNA accessibility measurements, and performed bi-modal UMAP dimensionality reduction, where we observed a clear separation of two major clusters - ECs and mural cells (Fig. 2B). A peak-to-gene linkage analysis of cell-type specific genes, such as COL5A1 and PDGFB (Fig. 2C), further validated the clear cell type separation at this time point (Fig. S4B).

**Figure 2.**
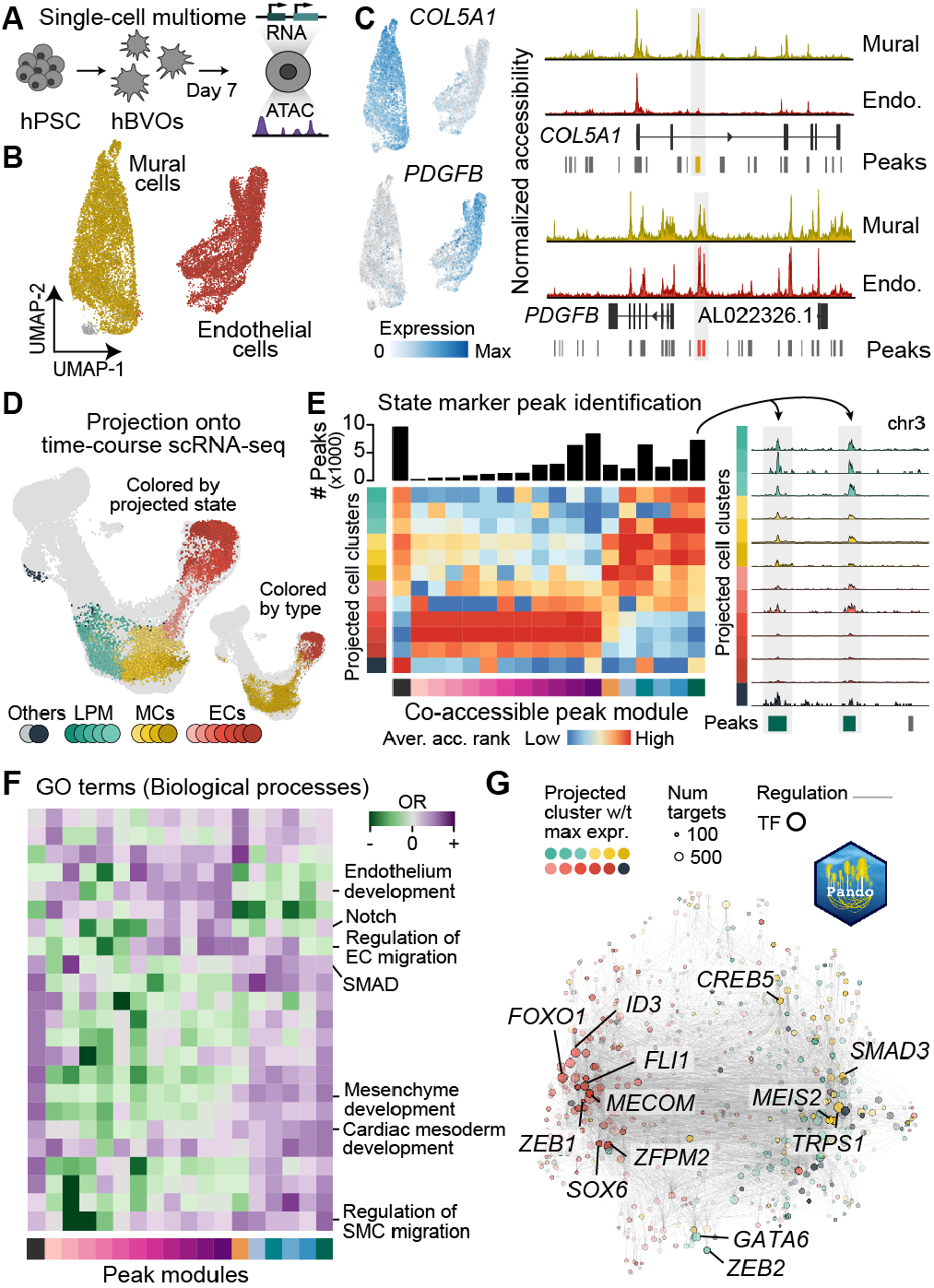
Paired single-cell RNA and chromatin accessibility profiling reveals candidate regulators of endothelial and mural cell fate maintenance. (A) Schematic of scMultiome (paired scRNA-seq and scATAC-seq) experiment performed at day 7 of hBVO development. hPSC, human pluripotent stem cells. (B) A WNN-based UMAP embedding of scRNA-seq and scATAC-seq with cells colored by cell type. Insert in the top left is colored by cell line. MC, mural cells; EC, endothelial cells. (C) Feature plots and coverage plots of COL5A1, marking mural cells, and PDGFB, marking endothelial cells. (D) Gene expression profile of day 7 hBVO scMultiome data projected to the scRNA-seq time course data with cells colored by cell cluster based on analysis in Fig. 1E. Insert in bottom right presents cells colored by cell type based on analysis in (B). (E) Normalized accessibility of example genes belonging to the same co-accessible peak module across projected cell type clusters from the time course scRNA-seq data. Arrows point to an example of a differentially accessible peak across cell type clusters. Aver. acc. rank, average accessibility rank; Norm. access., normalized accessibility. (F) Heatmap of enrichment or depletion of chosen gene ontology (GO) biological processes in co-accessible peak modules based on GREAT enrichment analysis. (G) UMAP embedding of the inferred TF network based on co-expression and interaction strength between TFs computed with Pando GRN framework. Circles: transcription factors (TFs) colored by projected cell type cluster. Circle size: difference in target numbers. See also Fig. S4.

Next, in order to perform cell type sub-clustering at a finer resolution, we projected the gene expression data obtained in the scMultiome experiment to the time course transcriptomic atlas data we had generated previously (Fig. 2D; Fig. S4C). We then transferred cluster labels from the time course data to the scMultiome data using a k-nearest neighbor (kNN) classifier (He et al., 2020). As expected, the projected cells mapped to ECs and mural cells, as well as to endothelial progenitor cells bridging the two cell type populations. Next, we identified peaks showing significant accessibility differences among different cell clusters and ranked the accessibilities of all clusters for each cluster-variable peak to obtain co-accessible peak modules (Fig. 2E; Fig. S4D, E; Table S4). To determine the distinct biological functions associated with the chromatin accessibility profile of each co-accessible peak module, we conducted Genomic Regions Enrichment of Annotations Tool (GREAT) analysis (Mao et al., 2012; McLean et al., 2010) (Fig. 2F). GREAT defines a regulatory domain for each gene and thus associates genomic regions with genes, followed by statistical analysis of ontology enrichments of associated genes. We found mural cells to be strongly associated with cardiac mesoderm and mesenchyme development, as well as with regulation of SMC migration. Furthermore, we observed an enrichment in gene regions associated with Smad protein signal transduction, which has been previously linked to smooth muscle cell development, proliferation and migration in primary human aortic SMCs (Tang et al., 2011) and in a mouse model (Mao et al., 2012). Endothelial cell clusters were associated with regulation of endothelium development, with some cell states being highly associated with Notch signaling and blood vessel EC migration.

We then applied the Pando gene regulatory network (GRN) framework (Fleck et al., 2021) to infer regulatory relationships between TFs, their target genes and regulatory genomic regions with the purpose to pinpoint molecular drivers of endothelial and mural cell identity maintenance and maturation (Fig. 2G; Table S5). Pando incorporates scATAC-seq data to identify conserved non-exonic regions and candidate accessible cis-regulatory elements and uses a linear model to infer directional regulatory relationships. We visualized the GRN using a UMAP embedding, which revealed groups of TFs that are involved in early mesoderm differentiation events, such as ZEB2, shown to be important in embryonic hematopoiesis and epithelial-to-mesenchymal transition (Goossens et al., 2011; Vandewalle et al., 2009) and GATA6, previously reported to instruct SMC development (Losa et al., 2017) and to play a role in EC function and survival (Froese et al., 2011). TFs leading to the formation of ECs were FLI1, MECOM, FOXO1, ZEB1 and ZFPM2. FLI1 has been shown to repress myogenic and promote endothelial differentiation (Ferdous et al., 2021). ZFPM2 (a member of the Friend of Gata family) is crucial for the maintenance of the coronary vasculature in mice (Fog2) (Zhou et al., 2009), while FOXO1 has been identified as an essential regulator of EC growth which promotes EC quiescence by antagonizing MYC and decelerating metabolic activity (Wilhelm et al., 2016) associated with EC maturation (Zecchin et al., 2017). MECOM, and particularly its isoform EVI1, have been shown to play a major role in hematopoietic stem cell differentiation and Evi1-/-mouse embryos fail to form an organized vascular network (Yuasa et al., 2005) and Evi1^δex3/δex3^ mouse embryos exhibit congenital heart defects (Bard-Chapeau et al., 2014). Interestingly, both ZEB1 and ZEB2 have been reported as inducers of endothelial-to-mesenchymal transition - a cellular transdifferentiation mechanism involved in developmental (e.g. cardiogenesis) and pathological (e.g. tumor metastasis) processes (Kovacic et al., 2019; Piera-Velazquez and Jimenez, 2019). SMAD3 and TRPS1 positively regulated the acquisition of mural cell fate in agreement with previous reports of the importance of these TFs for cranial and cardiac neural crest cell (Gong et al., 2020; Tsai et al., 2009) and SMC development (Machon et al., 2015), respectively. Altogether, these data show that regulatory region accessibility and TF expression reflect transitions of organoid cell development and cell type specification in accord with previous reports on blood vessel development.

### *In organoid* perturbations decipher endothelial versus mural cell specification

To investigate the mechanisms regulating hBVO development and in particular the bifurcation into mural and ECs, we leveraged a pooled genomic perturbation experiment with single-cell transcriptome readout (CROP-seq) (Datlinger et al., 2017) (Fig. 3A). We designed gRNAs targeting 16 receptors and 18 TFs (3 gRNA per gene) (Fig. S5A), which were differentially expressed throughout hBVO development (Fig. 3B). We transfected iCas9 hiPSCs with two different pooled gRNA lentiviral libraries, each containing gRNAs targeting a total of 17 genes and a set of 8 non-targeting gRNAs as a control with a vector expressing GFP as a selection marker (Fig. S5B). Next, we sorted vector positive cells based on green fluorescent protein (GFP) expression and induced Cas9 expression by doxycycline treatment for 4-6 days (Fig. S5C). We then generated mosaic hBVOs containing a multitude of wild-type (WT) and knock-out (KO) genotypes (see Methods).

**Figure 3.**
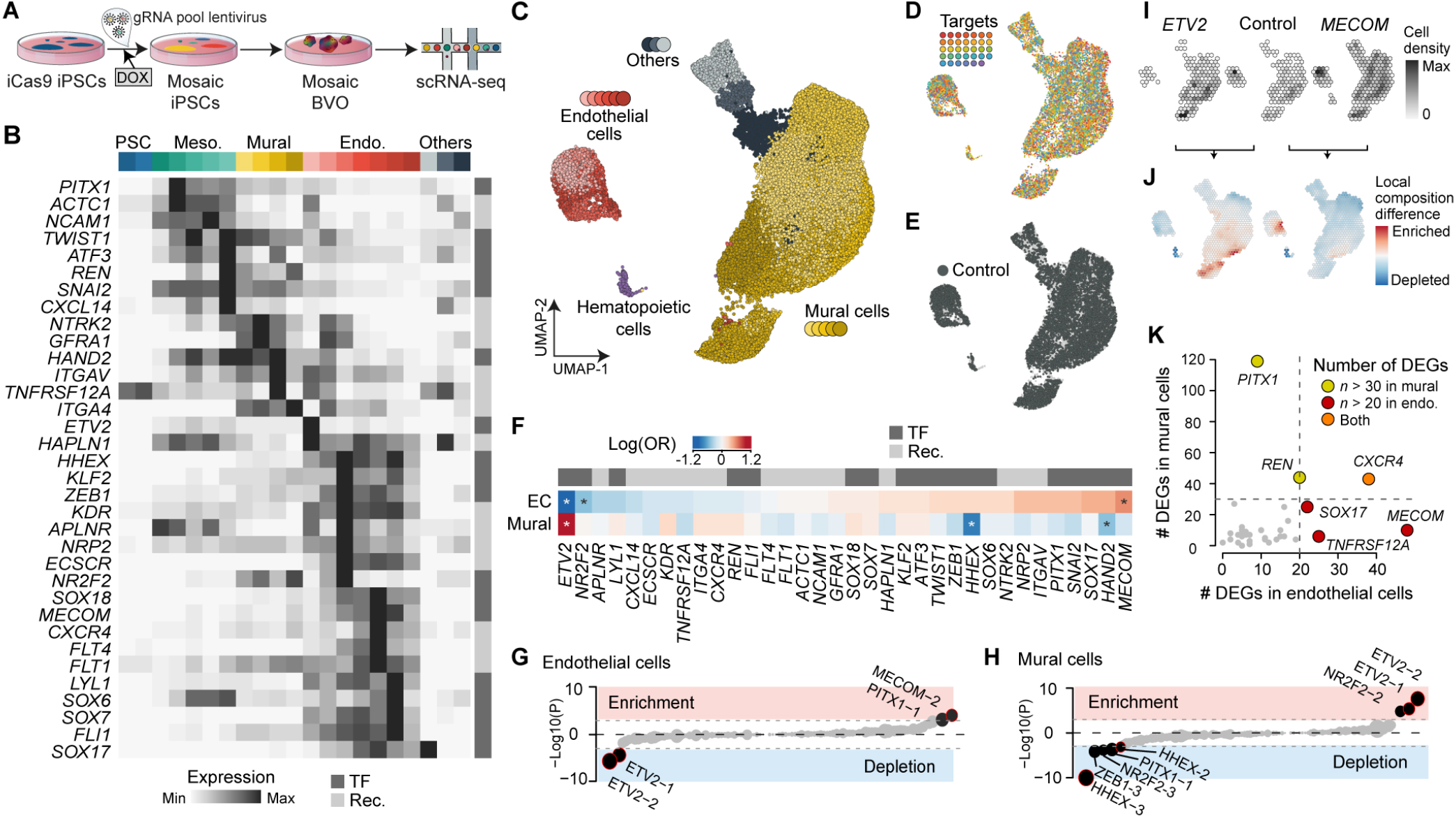
Multiplexed *in organoid* CRISPR perturbations can decipher endothelial versus mural cell specification. (A) Schematic of single-cell perturbation in hBVOs using the CROP-seq method. A doxycycline inducible Cas9 line (iCRISPRn) transfected with pooled gRNA libraries targeting a total of 16 receptors and 18 TFs (3 gRNA/gene), and containing 8 non-targeting gRNAs. Cas9 was induced by the addition of Doxycycline (Dox) for 4-6 days and used for the generation of hBVOs. (B) Heatmap showing average gene expression of targeted receptors and TFs in cell type clusters of developing hBVOs. Rec., receptor; TF, transcription factor. (C - E) UMAP embedding with cells colored by cell type (C), by detected targeting gRNA (D), or non-targeting gRNA (E). (F) Heatmap showing the effect of gRNAs (average of all 3 gRNAs/gene) on the proportion of mural or endothelial cells, represented as the log odds ratio. Significant differences (BH-corrected P<0.1) are indicated with a star. (G, H) Lollipop plot showing the impact of individual gRNAs on the proportion of endothelial (G) and mural (H) cells. (I) UMAP embedding colored by cell density with nearby cells summarized together in hexagonal bins. Cell den., cell density. (J) UMAP embedding colored by enrichment/depletion of gRNAs targeting ETV2 or MECOM relative to control (dummy gRNAs) with nearby cells summarized together in hexagonal bins. (K) Scatter plot of number of differentially expressed genes (DEGs) in mural, endothelial cells, or both in cells in which a gRNA has been detected. See also Fig. S5.

At days 7, 14 and 21 of differentiation, we sequenced single-cell transcriptomes and gRNA complementary DNA (cDNA) amplicons and recovered 35,639 cells, in 21,244 of which at least one gRNA was detected (Fig. S5D, E; Table S1). We integrated all cells using CSS, and generated a UMAP embedding where cells were clustered and annotated based on marker gene expression (Fig. 3C; Fig. S5F, G). In addition to ECs (CLDN5^+^, PECAM1^+^) and a large number of mural cells (PDGFRB^+^, COL1A2^+^), we detected the presence of hematopoietic-like cells (PTPRC^+^) and some offtarget cells (PAX3^+^ or EPCAM^+^) (Fig. S5G). Importantly, targeting and non-targeting gRNAs were detected across all cell types (Fig. 3D, E) and gene expression of the majority of target genes was downregulated (Fig. S5H).

To determine the effect of gene perturbation on cell fate, we tested the association of single gRNA detection with major cell type abundance (Fig. 3F; Fig. S5I). ETV2 gRNA detection was significantly associated with depletion of ECs and enrichment of mural cells, confirming the essential role of ETV2 for EC development in hBVOs (Kataoka et al., 2011; Lee et al., 2008). Another gRNA linked with a significant negative effect on EC differentiation was targeting NR2F2, a gene strongly associated with venous EC fate which is also expressed in mesenchymal cells during organogenesis and whose KO in mice leads to vascular abnormalities in the heart and brain in utero (Pereira et al., 1999). Remarkably, gRNAs targeting MECOM, a gene expressed highly in ECs of developing hBVOs (Fig. 3B), were enriched in ECs and depleted in a large proportion of mural cells (Fig. 3F-K; Fig. S5I-L). We identified differentially expressed genes (DEGs) between control and MECOM-KO cells and uncovered upregulation of THRB, IGFBP5, NRP2 and NEFH in MECOM-KO ECs (Fig. S5L). Furthermore, we observed a strong depletion of MECOM-and ETV2-gRNAs in hematopoietic cells. MECOM isoforms have an established role in the promotion of leukemogenesis (Birdwell et al., 2021), but are also involved in mesenchyme development (Bard-Chapeau et al., 2014; Hoyt et al., 1997) and are required for endothelial-to-hematopoietic transition of aortic endothelium, referred to as hemogenic endothelium, a Notch-dependent process (Goyama et al., 2008; Konantz et al., 2016). These findings about the role of MECOM in development in combination with the results of enrichment-depletion analysis suggest that, while MECOM is upregulated in ECs, it is not required for EC development and is instructive for the differentiation of hematopoietic cells and subtypes of mural cells in hBVOs.

In contrast to MECOM, we observed an enrichment of gRNAs targeting HHEX in hematopoietic-like cells (Fig. S5I), as well as depletion in mural cells (Fig. 3F, H). Among other functions, HHEX serves as a negative regulator of hematopoietic and non-lymphatic endothelium development (Gauvrit et al., 2018; Hallaq et al., 2004; Kubo et al., 2005). Nevertheless, in a study led by Hallaq et al, a null mutation of Hhex in mice resulted in abnormal vasculogenesis and the authors suggested this could be due to the depleted or absent SMCs they observed (Hallaq et al., 2004). Our approach corroborates these findings in the context of human development.

Among the other genes whose KO decreased mural and favored EC differentiation were HAND2, SNAI2, PITX1 and NRP2 (Fig. S5I, J). HAND2, SNAI2, and NRP2 are expressed in the developing mesenchyme and necessary for SMCs maturation (Yamagishi et al., 2000) and proliferation (Coll-Bonfill et al., 2016; Pellet-Many et al., 2015). KO of PITX1, a gene associated with cartilage, bone and muscle hindlimb development (Marcil et al., 2003; Wang et al., 2018), resulted in decreased mural cell fate acquisition and a large number of DEGs in mural cells (Fig. 3F, H, K; Fig. S5I, J). Altogether, these data provide support for previous findings about vasculature development, highlight the capability of hBVOs to mimic vascular development and open new avenues for exploration.

### Transplantation of human BVOs into immunocompromised mice facilitates endothelial arteriovenous differentiation

BVOs have been previously suggested to further mature after perfusion resulting from transplantation into immunodeficient mice (Wimmer et al., 2019b). To explore the transition to mature vessels, we set out to explore the effect of transplantation on EC maturation and specification. To this aim, we transplanted day 14 hBVOs under the kidney capsule of immunocompromised mice (Fig. 4A). Immunohistochemical analysis confirmed the presence of vascular networks and blood vessels formed by ECs, covered by a basement membrane (Fig. 4B). Two months after transplantation, we sacrificed the mice and collected hBVOs. After dissociation, we stained all cells against human CD31 (hCD31) and hPDGFR-β, marking human endothelial and mural cells, respectively, FACS sorted marker positive cells (Fig. S6A) and performed single-cell transcriptomics. We annotated cells (16,410 human cells) based on marker gene expression and comparison to a single-cell transcriptomic atlas of the human developing gut (Yu et al., 2021) (Fig. S6B, C, D; Table S1).

**Figure 4.**
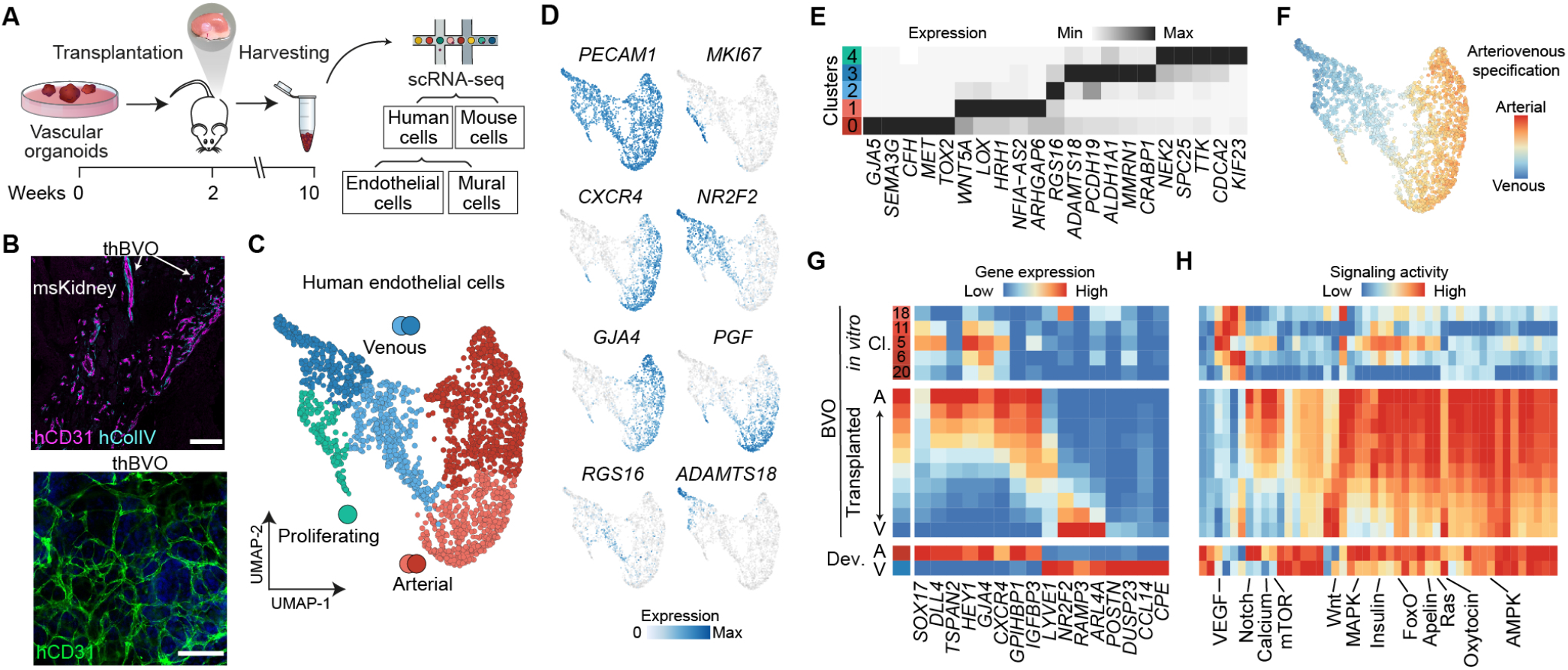
Transplantation of hBVOs into immunocompromised mice facilitates endothelial arteriovenous differentiation. (A) Schematic of day 14 hBVO transplantation into immunocompromised mice, followed by organoids isolation, human cell sorting (hCD31^+^, endothelial and PDGFR-β^+^, mural) and scRNA-seq 2 months later. (B) Immunohistochemical characterization of transplanted hBVOs (thBVOs). msKidney, mouse kidney. Scale bars: 200 µm (top) and 100 µm (bottom). (C) UMAP embedding of human endothelial cells (hCD31^+^) with cells colored by cell subtype based on marker gene expression. (D) Feature plots colored by expression of cell type marker genes. (E) Heatmap showing average expression of cluster marker genes in ECs from thBVOs. (F) UMAP embedding colored by arteriovenous (AV) cell specificity. (G, H) Heatmap of gene expression of artery-or vein-enriched genes (G) and signaling pathway activation (H) across endothelial cells in *in vitro* and transplanted hBVOs and in the developing human gut tube. A, arterial; V, venous; Dev., developing human gut tube. See also Fig. S6.

There was a surprising abundance of PDGFRB-positive mesenchymal cells that became prevalent in transplanted hBVOs (thBVOs) (Fig. S6E, F). The majority of these mesenchymal cells strongly resembled pericytes (PDGFRB-high), SMCs (ACTA2-high) and fibroblasts (LUM-high), while more rare cell populations were phenotypically similar to adipocytes (CDF-high) and chondrocytes (MATN4-high) (Fig. S6B, C, E, F)). We analyzed ECs in the transplanted hBVOs and identified three main EC populations - arterial-like (CXCR4-high), venous-like (NR2F2-high), and proliferating ECs (MKI67-high) (Fig. 4C, D). Arterial ECs could be further divided into two subpopulations: cluster 1 (c1) was marked by the expression of canonical arterial EC markers such as connexins GJA4 and GJA5, and the semaphorin SEMA3G, while cluster 0 (c0) showed upregulation of PGF, ARHGAP6, WNT5A, and HRH1 (Fig. 4E). PGF and ARHGAP6 have been previously associated with tip (Goveia et al., 2020) and capillary (Schupp et al., 2021) ECs, respectively, while WNT5A (Skaria et al., 2017) and HRH1 (Ashina et al., 2015) have been shown to positively regulate EC permeability (Fig. 4D, E). Venous-like cells could be subdivided into two clusters: cluster 3 (c3) was enriched in the metalloproteinase ADAMTS18 (Sissaoui et al., 2020), the adhesion protein MMRN1 (Sabbagh et al., 2018), members of the retinoic acid signaling pathway (CRABP1 and ALDH1A1) (Napoli, 2016), and genes involved in adhesion and barrier formation (PCDH19) (Higurashi et al,. 2015), whereas cluster 2 (c2) had increased expression of RGS16, a gene involved in glucose homeostasis (Villasenor et al., 2010).

We generated an arteriovenous-specificity score based on a previously published primary human developmental EC dataset (Yu et al., 2021) and assigned a score to each cell, further confirming the arteriovenous specificity of different clusters (Fig. 4F). We identified genes with distinct expression in primary human arterial and venous ECs, respectively, and analyzed their expression in EC clusters of *in vitro* and transplanted hBVOs (Fig. 4G). Interestingly, *in vitro* hBVO ECs had an overall higher expression of arterial EC markers, such as SOX17, DLL4 and HEY1 with only one cluster (c18, derived from day 6, 7 and 14) showing increased expression of venous cell markers such as the TF NR2F2, as well as POSTN and ARL4A. The latter two genes, however, are also expressed in mural cells suggesting that this subpopulation constitutes an immature venous-like state or an intermediate population of cells between mural and endothelial cell lineages. On the other hand, ECs from thBVOs displayed clear arterial and venous states, marked by increased expression of DLL4 and CXCR4, and NR2F2 and ARL4A, respectively.

Signaling activity analysis revealed a strong impact of transplantation on cell-cell communication within hBVOs (Fig. 4H, Fig. S6F). Pathways involved in hormone (Relaxin, Glucagon, Estrogen) and immune (T cell receptor, B cell receptor) signaling showed an increased signaling activity in hBVO ECs 2 months after transplantation, with scores comparable to developing human ECs. Interestingly, we noticed that transplantation caused the inhibition of several pathways in hBVO ECs, such as VEGF, ErbB and TGF-β, which correlated with a decreased activity of these pathways *in vivo*. We also detected the presence of signaling pathways (Hippo, JAK-STAT and Hedgehog), which were less active in hBVO ECs in both *in vitro* and *in vivo* context compared to primary ECs. This suggests that additional cues - growth factors, immune cells as well as missing hemodynamic forces, such as shear stress, which affect endothelial mechanotransduction - are required for maturation and further differentiation of hBVO ECs *in vitro*. Furthermore, our signaling analysis highlighted the resemblance of thBVO ECs to primary arterial and venous ECs, underscoring the particular degree of plasticity which hBVO ECs exhibit prior to transplantation. Both *in vivo* and primary human venous ECs had notably higher Wnt and Oxytocin signaling activity scores in comparison to arterial ECs. Arterial ECs on the other hand were enriched in transcripts important in Notch, VEGF and Ras signaling, in accord with previous findings about the importance of these pathways in arteriovenous specification in zebrafish (Lawson et al., 2001, 2002; Ren et al., 2013). Altogether, these results suggest that transplantation of hBVOs markedly increases their similarity to *in vivo* blood vessel ECs and induces arteriovenous differentiation, while mural cells retain multipotent properties.

### Manipulation of VEGF and Notch pathways in developing BVOs impacts EC differentiation and shifts arteriovenous-like endothelial cell states

We next wanted to utilize the hBVO system to explore how developing human vasculature responds to perturbation of key signaling pathways. Notch and VEGF signaling pathways, highlighted in our analysis of thBVOs, are well-known regulators of EC differentiation, proliferation, angiogenesis and arteriovenous specification (Akil et al., 2021; Casie Chetty et al., 2017; 2006), and numerous studies have shown that their effect on vascular cells is highly context specific and dose-dependent (Cao et al., 2009; Luo et al., 2021; Pontes-Quero et al., 2019a). We therefore performed a screen of 22 different conditions where we pharmacologically perturbed VEGF and Notch pathways in hBVOs in an *in vitro* context and measured their effect on the expression of multiple endothelial and mural cell markers using qPCR (Table S7) (Fig. S7A-C). We selected 5 of these conditions for further characterization based on their effect on relative EC abundance and arteriovenous specification including different concentrations of VEGF-A (100 ng/ml, VEGF-A^high^; 50 ng/m, VEGF-A^mid^; 10 ng/ml, VEGF-A^low^) and gamma-secretase inhibitor DAPT (Notch pathway inhibitor, Notch-inh; 5 µM) added between day 5 and 10 of hBVO development (Fig. 5A). These perturbations had a distinctive impact on network sprouting and proliferation which could be observed by brightfield microscopy (Fig. 5B). Immunostaining against the EC markers CD31 (adhesion molecule) and TIE1 (tyrosine kinase) revealed decreased endothelial cell-cell interaction and vessel formation upon Notch inhibition (Fig. 5C). In VEGF^mid^ and VEGF^low^ conditions, we observed a large number of APLNR^+^ ECs with low CD31 expression sprouting from the networks and leading the angiogenic front.

**Figure 5.**
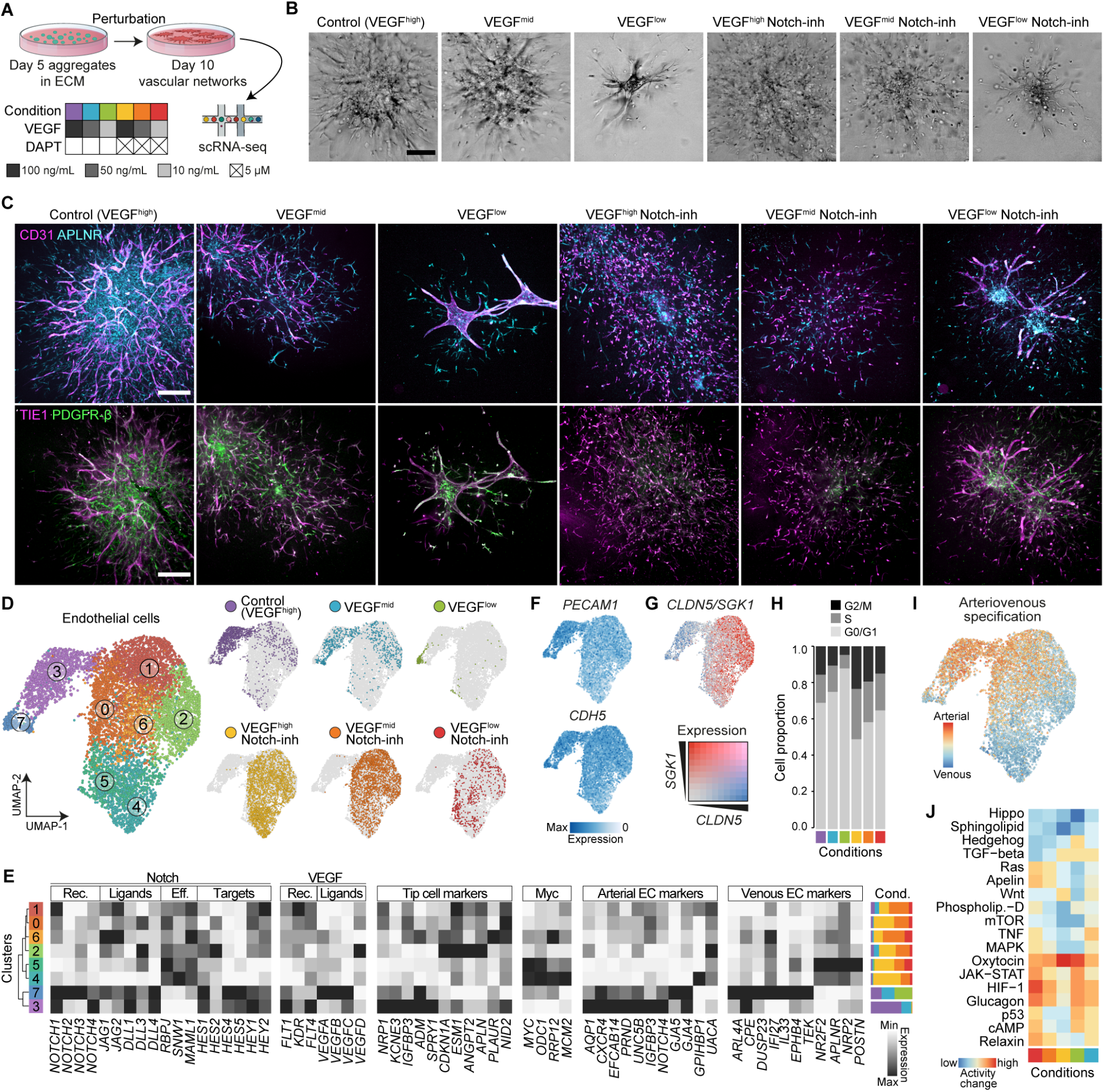
Manipulation of VEGF and Notch pathways in developing BVOs impacts EC differentiation and shifts arterial-venous-like endothelial cell fate proportions. (A) Schematic of experimental setup for the perturbation of signaling conditions. Mesoderm aggregates were embedded in extracellular matrix (ECM) at day 5 and treated following the original BVO protocol, or with a signaling perturbation (decreased VEGF or addition of DAPT, a Notch-inhibitor). Samples were dissociated and scRNA-seq was performed at day 10. (B) Brightfield images of hBVOs treated with different signaling perturbations. Scale bar: 200 µm. (C) Immunohistochemical staining detecting CD31 (magenta) and APLNR (cyan) (top row) or TIE-1 (magenta) and PDGFR-β (green) (bottom row) in cells across control and pharmacologically perturbed hBVOs at day 10 of differentiation. Scale bar: 250 µm. (D) UMAP embedding of endothelial cells (ECs) colored by cluster (left) as well as colored and split by signaling pathway perturbation (right). (E) Feature plots of the endothelial adhesion cell markers PECAM1 and CDH5. (F) Co-expression feature plots of the endothelial adhesion cell marker CLDN5 and cell migration-associated SGK1. (G) Proportional stacked bar graph of proportion of cells in each cell cycle phase, stacked by condition. Cond., condition. (H) Heatmap of average gene expression of Notch ligand, receptor, effector and target genes, VEGF ligands and receptor genes, tip-cell marker genes, genes involved in Myc signaling, arterial- or venous-specific EC marker genes across EC clusters. (I) UMAP embedding of ECs colored by arteriovenous specificity, defined as the arithmetic difference between the arterial and venous signature scores. (J) Heatmap of signaling pathway activity across signaling pathway perturbation normalized to control cells from ECs only. See also Fig. S7.

We performed scRNA-seq of organoids from these 5 conditions and control on day 10 and recovered 51,261 cells (Fig. S7D; Table S1), which we could annotate as ECs and mural cells based on marker gene expression (Fig. S7E, F). We subsetted ECs, performed unsupervised clustering resulting in 8 distinct clusters (c0-7), and projected the data into two dimensions using UMAP. The embedding illuminated alterations in molecular EC states induced by the signaling perturbations, with c3 being enriched in control, c7 being enriched in VEGF^low^ condition, and all remaining clusters being strongly enriched in Notch inhibition (DAPT treatment) conditions, independent of the level of VEGF (Fig. 5D, Fig. S7L). In all clusters, we assessed expression of marker genes as well as of a curated set of genes involved in Notch signaling, VEGF signaling, tip-cell identity, Myc signaling, as well as arteriovenous specification (Fig. 5E). Notch inhibition generally led to a strong downregulation of Notch ligands (DLL1, DLL3, DLL4), receptors (NOTCH1, NOTCH2, NOTCH3, NOTCH4) and target genes (HES1, HES2, HES4, HES5) and an upregulation of Notch effector genes (RBPJ, repressor; MAML1-3, EP300, co-activators) (Fig. 5E). Interestingly, this change was specific to ECs and could not be detected in mural cells upon Notch inhibition (Fig. S7G). Furthermore, ECs of organoids treated with DAPT induced a lower average expression of typical endothelial cell-cell adhesion markers such as CLDN5, PECAM1 (Fig. 5F) (Herbert and Stainier, 2011), and CDH5, and an increased SGK1 (Fig. 5G) expression, which is associated with increased permeability (Claesson-Welsh et al., 2021) as well as with cell motility (Abraham et al., 2009; Sang et al., 2020). This supports our imaging-based observation that Notch inhibition leads to increased hBVO vessel sprouting.

Interestingly, a combination of Notch inhibition and VEGF^high^ condition led to a strong increase of S phase and G2/M phase of the cell cycle in ECs, but not in mural cells (Fig. 5H, Fig. S7H, I, N). The induction of EC proliferation resulted in an increased number of ECs versus mural cells under Notch-inhibition and in VEGF^high^ conditions, with additive effects of the two variables. These results supported the immunohistochemical analysis of hBVOs showing a decreased number of mural cells (PDGFR-β^+^) in DAPT-treated organoids (Fig. 5C) and are in alignment with previous findings that low levels of Notch and high levels of VEGF signaling promote EC proliferation (Phng and Gerhardt, 2009).

Proliferating ECs were enriched in c4, which was marked by expression of MKI67, MYC (a direct target of Notch), its downstream genes, and NR2F2, which have been associated with inhibition of arterial and induction of venous cell fate (Fig. 5E, Fig. S7L) (Luo et al., 2021; Sissaoui et al., 2020; Su et al., 2018). In addition, this cluster showed lowest expression of genes previously shown to be upregulated in tip cells (DLL4, ESM1, ANGPT2, CXCR4, APLN, KCNE3, IGFBP3, ADM, NID2, SPRY1, PLAUR) (Fig. 5E). Tip/stalk cell specification, arteriovenous differentiation and EC motility have all been previously associated with high ERK activation and temporary cell cycle arrest (Luo et al., 2021; Mühleder et al., 2021; Yasuda et al., 2016), highlighting the importance of the cell cycle for EC differentiation and specification. Notably, tip cell markers were heterogeneously expressed in the other EC clusters, in particular in clusters 0, 1, 2, 6 and 3 enriched in VEGF^high^ and VEGF^mid^ (no Notch inhibition) conditions, suggesting phenotypic heterogeneity of tip cells affected by differences in signaling cues - a question which has only recently been addressed (Zarkada et al., 2021). C2 showed high expression of ESM1, ANGPT2 and APLN as well as CDKN1A (encoding p21) transcripts. p21 is a cell cycle inhibitor which has been identified as an effector of an ERK-dependent hyper mitogenic cell cycle arrest mechanism responsible for low tip cell proliferation (Pontes-Quero et al., 2019b).

To explore arteriovenous specification within the hBVOs *in vitro*, we assigned each EC an arterial and venous score based on arterial and venous EC gene expression from a published scRNA-seq dataset of primary human ECs in the developing gut tube (Yu et al., 2021) (Fig. 5I). Based on these analyses and supported by the expression of known arterial and arteriole EC marker genes (AQP1, CXCR4, IGFBP3, GJA4), we observed that cells from c3 most strongly resemble arterial cells (Fig. 5E, I). This cluster was enriched for cells from VEGF^high^ (control) and VEGF^mid^ conditions without Notch inhibition. On the other hand, venous-like cells were found in c5 and c7 based on an elevated venous score and supported by the expression of vein and venule EC marker gene expression (NR2F2, APLNR, POSTN in c5; ARL4A, CPE, DUSP23, TFPI in c7) (Fig. 5E). Interestingly, each of the two venous-like cell populations expressed only a subset of venous marker genes and they were found across different signaling perturbation conditions. C5 was composed of cells from Notch inhibition conditions independent of the level of VEGF. In contrast, c7 contained all cells from VEGF^low^-treated hBVOs, consistent with reports that low VEGF levels are required to promote and maintain vascular EC differentiation and survival, while high VEGF levels induce arterial and inhibit venous EC specification (Casie Chetty et al., 2017; Zhang et al., 2008).

Next, we assessed how signaling is generally affected by our VEGF and Notch perturbations and therefore quantified the activity of signaling pathways in hBVO ECs across signaling perturbation conditions. This was done by subsetting the single-cell expression data to genes associated with signaling pathways based on the KEGG database (excluding ligands), and calculating SCENIC-inferred regulome activities of TFs involved in each given signaling pathway (Fig. 5G; Fig. S7K; Table S5). Remarkably, we observed that VEGF^low^ conditions had a divergent effect on the activation of multiple major signaling pathways (Ras, Apelin, Relaxin, TNF) dependent on whether or not Notch was inhibited at the same time, pointing towards a complex and fine balance between these two signaling pathways. Signaling pathway activity in mural cells exhibited a similar pattern with a large increase of multiple signaling scores upon Notch inhibition (Fig. S7R). We observed increased signaling activity among pathways which have been associated with the promotion of VEGF expression and angiogenesis - JAK-STAT (Auzenne et al., 2012; Xue et al., 2017), cAMP (Amano et al., 2001), HIF-1 (Manalo et al., 2005), and Glucagon (Wang et al., 2014), - suggesting that such signaling activation might be a compensatory mechanism ensuring the maintenance of vascular cell homeostasis under reduced external VEGF supply.

While supportive mural cells have been implicated in angiogenesis and blood vessel maintenance, the understanding about their role in modulating the effect of VEGF, Notch, and other relevant cues on ECs remains limited. We discovered that mural cell populations (c5, Fig. S7P) expressing canonical pericyte / SMC markers such as PDGFA and MCAM, an essential regulator of pericyte-EC interaction (Chen et al., 2018), were reduced upon a decrease in VEGF concentration and were not present in organoids treated with the Notch inhibitor (Fig. S7M, O-Q). Furthermore, mural cells of organoids in Notch inhibition condition had a lower number of cells expressing the markers of mesenchymal differentiation PPRX1 and TWIST1 (c4) in comparison to non-treated organoids. Interestingly, Notch inhibition, particularly in combination with low VEGF concentration, led to an increase in the expression of CTGF, ANKRD1 and CYR62, YAP/TAZ target genes which have been associated with tumorigenic phenotype (Li et al., 2019; Marti et al., 2015). Collectively, these data support the notion that Notch and VEGF signaling affect the differentiation of vascular cells in an interdependent manner and hBVOs are a suitable model for studies of signaling pathways in vascular development.

### Assessment of human BVOs to model human vascular disorders

ECs across organs display unique molecular profiles (Nolan et al., 2013) and their heterogeneity is often predicated by vascular bed-specific and organ-specific disease manifestation (Potente and Mäkinen, 2017). In order to assess organ specificity of hBVO ECs, as well as their potential for disease modeling and regenerative medicine, we compared hBVO ECs to ECs derived from 14 different organs in a published single-cell transcriptomic dataset of human development (Cao et al., 2020) (Fig. 6A). We calculated the transcriptomic similarity between ECs from *in vitro* or transplanted hBVOs and ECs from different human developing organs (Fig. 6B). The hBVO EC transcriptome profile (both *in vitro* and transplanted) was most similar to ECs in the developing human pancreas and to a lesser extent similar to ECs in developing human muscle, intestine and stomach. hBVO ECs were most dissimilar to ECs of the developing human liver. To further explore the potential of hBVOs to serve as proxies of organ specific ECs, we examined the expression of genes associated with various important biological processes including the gene ontology (GO) terms “Transmembrane transport”, “Response to stimuli”, Tissue morphogenesis” and “Cell-cell signaling” (Fig. 6C). Interestingly, we observed that cerebellum-and cerebrum-associated blood vessel ECs were highly enriched in genes associated with these processes, in a pattern distinct from ECs of the other organs in the dataset. This finding is consistent with the tight cell-cell adhesion and low permeability of the blood-brain barrier, which necessitate the presence of a large number of selective transporters for the proper nutrient and metabolite movement between blood and brain (Segarra et al., 2021). ECs from other organs, such as kidney and pancreas, shared similarities in gene expression patterns. Curiously, there was a notable heterogeneity of ECs derived from the same organ with most arterial cells showing upregulated expression of GJA4, lymphatic cells showing expression of PROX1, and a specific subpopulation of “HB+” cells in a subset of organs expressing genes instructing hemoglobin production (HBG1, HBG2, HBA1). These data illuminate tissue-specific endothelial cell signatures and highlights differences and similarities between hBVO ECs and subtypes of primary ECs throughout the human body.

**Figure 6.**
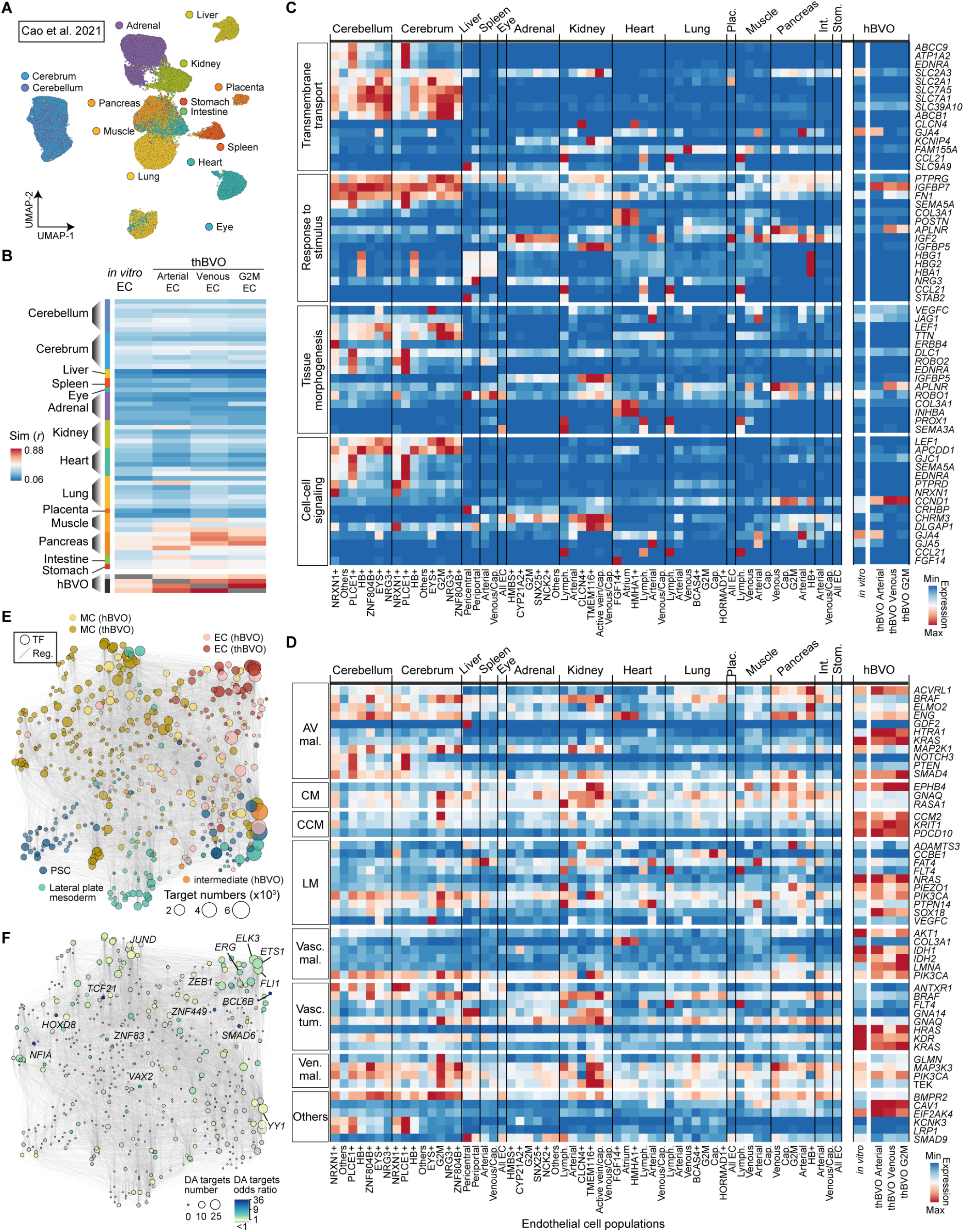
Assessment of human BVOs to model human vascular disorders. (A) UMAP embedding of primary endothelial cells from a reference human fetal gene expression atlas (Cao et al., 2020), colored by the 14 different organs it encompasses. (B) Heatmap representing the transcriptomic similarity between ECs in the *in vitro* (dark gray) or transplanted (light gray) hBVOs and fetal primary ECs. hBVO ECs most closely resemble ECs in the human pancreas. (C) Average gene expression of genes associated with the GO terms “Transmembrane transport”, “Response to stimuli”, “Tissue morphogenesis” and “Cell-cell signaling” across clusters of human primary and hBVO ECs. (D) Average gene expression of genes associated with vascular anomalies and other diseases across clusters of human primary and hBVO ECs. AV mal., arteriovenous malformation; CM, capillary malformation; CCM, cerebral cavernous malformation; LM, lymphatic malformation; Vasc. mal., vascular malformations associated with other anomalies or overgrowth; Vasc. tum., vascular tumors; Ven. mal., venous malformation. (E) UMAP embedding of the inferred GRN based on co-expression and interaction strength between TFs computed with Pando. Circles represent transcription factors (TFs) and are colored by cell type. Circle size indicates difference in target gene numbers. PSC, pluripotent stem cell; MC, mural cell; EC, endothelial cell; thBVO, transplanted hBVO. (F) GRN in (C) with TFs colored by the differentially accessible (DA) target genes odds ratio. Circle size indicates difference in DA target gene numbers.

Some of the clearest examples of diseases that affect specific vessel types and can manifest in an organ-specific manner are vascular anomalies, comprising vascular tumors and vascular malformations. We therefore compiled a list of genes with loss-or gain-of-function mutations that are associated with arteriovenous, capillary, cerebral cavernous, venous and lymphatic malformations, vascular malformations associated with other anomalies, vascular tumors and other vascular diseases (Table S8) and mapped these genes to *in vitro* and transplanted hBVO ECs, as well as to primary human fetal ECs (Fig. 6D). The mapped genes were differentially expressed across organs and most genes were detected in both primary and hBVO ECs. An interesting observation was that genes associated with cavernous capillary malformations, abnormally formed blood vessels in the central nervous system, exhibited higher expression in hBVO ECs than in ECs from the human fetal cerebellum or cerebrum, despite the dissimilarity of hBVOs to brain ECs. This discrepancy hints towards additional factors underlying the pathological mechanisms of vascular anomalies.

For better understanding the regulatory mechanisms driving the expression of vascular disease-and vascular anomaly-associated genes, we used *in vitro* and transplanted hBVO EC transcriptomes to infer GRNs (Fig. 6E; Table S4). Disease-associated genes were found to be regulated by TFs throughout hBVO development, and regulators of endothelial gene expression such as ETS1, ERG, FLI, ZEB1, ELK3 had a particularly strong effect (Fig. 6F). Altogether, these analyses suggest that hBVOs have potential for modeling human vascular disorders *in vitro* and bear a close transcriptomic resemblance to vasculature of the human fetal pancreas, but also reveal the need for further engineering of the hBVO system.

## Discussion

The vasculature is a heterogeneous organ promoting proper nutrient and gas exchange between the blood and tissues throughout the human body. Insufficient vascularization underlies ischemic disorders such as myocardial infarction and stroke, as well as tissue necrosis. Excessive and abnormal angiogenesis, on the other hand, can contribute to the pathogenesis of diabetic retinopathy, vascular malformations, cancer and many others. Human BVOs provide a new inroad into studying each of these areas. We have presented the first comprehensive single-cell genomics atlas of developing hBVOs and their perturbed states, thus providing an extensive resource for studying human blood vessel development and disease using this powerful *in vitro* system. Studying early human vascular development has been challenging due to inaccessible human embryonic tissues, and extrapolating results from other animal systems has become the gold standard, despite large costs, time investment and questionable ethics (Smith et al., 2018; Würbel, 2017). Our in depth single-cell transcriptomic and immunohistochemical characterization of developing hBVOs identified the presence of a lateral plate mesoderm progenitor population which further acquires endothelial or mural cell fate. ECs emerge at day 4 of differentiation, one day after supplementation with VEGF-A, and exhibit transient expression of ETV2 - a potent inducer of hemangiogenic program (Liu et al., 2015). Our transcriptomic analysis showed that by day 14 of differentiation hBVO cells resemble human endothelial and perivascular cells at the early developmental stage and highlighted differences and similarities of these cell types to their primary counterparts. We detected the emergence of both arterial-like and venous-like EC states at day 14 and 21 of hBVO development, however both states lack clear maturation. These results highlight the strong potential of hBVOs to recapitulate blood vessel development and to be used to study healthy and pathological vascular processes *in vitro*, especially with respect to early development.

GRN and lineage tracing analyses provided information on transcriptional regulators and progenitor capacities to drive endothelial and mural development and maintenance in hBVOs. We perturbed multiple TFs and receptors in hBVOs, which revealed effects of gene KO on endothelial, mural and hematopoietic cell fate specification and largely support previous findings in mice and zebrafish. We also establish single-cell sequencing methods that can be harnessed for systematic perturbation of BVO development using small molecules and other culture modulations. Perturbation of Notch and VEGF pathways had strong effects on hBVO development. Coupled with single-cell sequencing our results support previous studies in primary human and mouse embryonic stem cell-derived ECs and zebrafish (Chappell et al., 2013; Magnussen and Mills, 2021; Sörensen et al., 2009) showing that EC response to Notch inhibition and increasing VEGF concentration are involved in a negative feedback loop. This is also consistent with previous reports showing that higher Notch activity is associated with inhibition of MYC and cell cycle suppression, supporting arterial differentiation (Luo et al., 2021; Sissaoui et al., 2020; Su et al., 2018), while overexpression of COUP-TFII (encoded by NR2F2) in pre-arterial ECs induces cell-cycle gene expression and compromises arterial development (Sissaoui et al., 2020; Su et al., 2018). Future studies manipulating other pathways, dissecting mural-endothelial cell interaction, and combining single-cell sequencing with epigenome profiling will provide new insights into human vascular development, with implications for therapeutics. Indeed, current antiangiogenic (Ioannidou et al., 2021) and vascular normalization strategies (Magnussen and Mills, 2021) for cancer treatment will profit from a better understanding of the contextdependent effect of VEGF and Notch signaling on vessel formation in a human model system.

Previous studies have shown that upon transplantation, various organoid systems exhibit progressive maturation (Mansour et al., 2018; Spence et al., 2011; Takebe et al., 2014; Tran et al., 2019). Similarly, we showed that hBVO ECs can further specify and mature into arterial- and venous-like ECs, highlighting the potential of hBVO cells to mimic later developmental stages. Signaling pathway analysis uncovered an upregulation of many pathways upon transplantation into a host, highlighting both the predictive power of hBVOs, while also illuminating the limitation of the *in vitro* system. The finding that mural cells *in vivo* differentiate into various cell types with mesenchymal origin, corroborates the numerous reports of shared features between pericytes and mesenchymal stem cells (Cathery et al., 2018). This unrestricted differentiation has implications for hBVOs as a therapeutic agent, since strategies need to be developed to protect against off-target differentiation. It will be important to understand the potency of each cell population within BVOs, and how they can differentiate in various permissive environments. We note that a major limitation of the hBVO system is the requirement for transplantation for functional maturation. It remains elusive what currently limits maturation *in vitro*. Flow and mechanical forces would allow investigation of functional parameters such as permeability or immune-cell adhesion and extravasation under different conditions. Our work provides a reference foundation for future efforts aimed at human blood vessel maturation and circulatory implementation entirely *in vitro*.

Vascular SMCs and pericytes are implicated in the production of basement membrane, which promotes proper blood vessel wall assembly, stabilization, and quiescence under healthy conditions (Stratman et al., 2017). Many previous human stem cell-derived endothelial systems lack mural cells and associated morphological development. Here we reconstruct differentiation trajectories of pericyte states from mural progenitor cells. Pericytes subsequently interact directly with ECs within hBVOs *in vitro* and after transplantation. This close interaction places them at an ideal position to respond to injury or insult in tissue-appropriate ways. In the future, it is important to consider mural cells when studying blood vessels in the context of disease because the failure of crosstalk between perivascular and endothelial cells leads to the development of a multitude of of congenital and acquired diseases, such as vascular malformations, retinopathies, and cancer (Sweeney and Foldes, 2018).

The incorporation of blood vessels into engineered tissues is crucial for achieving biologically relevant tissue and organ substitutes in regenerative medicine as well as for the development of accurate three-dimensional *in vitro* models which measure up to the high complexity of human physiology. The microvascular cells within each organ are uniquely adapted to serve angiocrine functions, satisfy metabolic demands, and ensure the proper development of the particular organ, with each organ microvasculature exhibiting differences in permeability, angiogenic potential and metabolism (Augustin and Koh, 2017). It is therefore key to generate organotypic hBVOs that allow the study of organ-specific diseases, and possibly be applied for tissue-regenerative therapy. Our analysis revealed that *in vitro* hBVOs resemble most closely the human pancreas microvasculature from a transcriptomic perspective. A similarity to the pancreatic EC transcriptome was further strengthened upon transplantation of hBVOs despite the fact that the organoids were located under the kidney capsule. In addition to site-specific microenvironmental cues which lead to immediate transcriptional changes, it has been shown that epigenetic mechanisms in different organs are specified early during embryonic development and mediate differential gene expression profiles of ECs across tissues, particularly in macrovascular beds (Augustin and Koh, 2017; Nakato et al., 2019). Hence, in order to find appropriate culture conditions for organ-specific hBVOs, one needs to first decipher the major drivers of the differences between vascular cells of different organs. Epigenetic memory needs to be taken into consideration to improve approaches which have already been applied for certain tissues, such as co-culture of PSC-derived ECs with the desired tissue (Giacomelli et al., 2017; Helle et al., 2021; Sances et al., 2018), culture in tissue-conditioned media (Puech et al., 2018), addition of small molecules (Roudnicky et al., 2020a) or synergistic overexpression of transcription factors (Dailamy et al., 2021; Roudnicky et al., 2020b). We highlight the high degree of intraorgan heterogeneity displayed by ECs, which likely governs the difference in development and severity of vascular disorders. In summary, we present a detailed single-cell multiomic atlas of developing hBVOs, with a focus on cell fate and state transitions during development under healthy and perturbed conditions. Collectively, this resource lays the foundation for studies of development and disease of the human vasculature utilizing hBVOs as a model.

## Limitations of the study

hBVO generation involves cumbersome steps such as embedding in a matrix and extracting single organoids, which would need to be optimized, e.g. by implementing robotic technologies, in order to achieve higher throughput for translational research. While our *in organoid* perturbation experiment gives useful information on the effect of gene KO on vascular development in a high-throughput manner and allows for KO induction at any time point of differentiation, this approach suffers from several disadvantages. There is the possibility, inherent to CRISPR/Cas9 KO techniques, of off-target effects and the acquisition of adaptive mechanisms, and the genotype of each single mutant (homo-or heterozygous) is not known. The mosaic design also makes it difficult to assess cell autonomous and non-cell autonomous effects of KO.

## AUTHOR CONTRIBUTIONS

M.T.N., R.A.W., M.S. generated organoids used in this study. M.T.N. generated all single-cell transcriptomics and multiomics datasets. R.A.W. transplanted organoids into host animals, supported dissociation and did immunohistochemistry of transplanted organoids. J.M.P. provided guidance on and infrastructure support for BVO culture and transplantation. M.S., M.T.N. generated iTracer organoids. M.T.N., M.S. generated CROP-seq libraries and transfected cells. M.T.N., J.M.N. performed immunohistochemistry and microscopy of all *in vitro* hBVOs. M.T.N., M.S. performed RT-qPCR of pharmacologically perturbed organoids and M.T.N. analysed the results. M.T.N., Z.H. performed the analysis of the scRNA-seq developmental time course, transplanted organoids, pharmacologically perturbed organoids. Z.H. performed analysis of lineage recording and primary endothelial cell datasets. Z.H. analysed CROP-seq and multiomics data with support from M.T.N.. M.T.N., Z.H., J.G.C., B.T. designed the study. M.T.N. drafted the manuscript with support from J.G.C., B.T., Z.H.. All authors edited and approved the manuscript.

## ACKNOWLEDGEMENTS

We thank the Treutlein and Camp labs for helpful discussion. We thank Sophie Jansen, Jonas Fleck and Giovanna Brancati for sharing experience, CROP-seq plasmids, primers and R scripts for pooled perturbation genomic experiments. We thank Ashley Maynard for discussion, sharing plasmids and primers for lineage recording experiments. We thank Rene Holtackers, Philipp Wahle and Ryo Okuda for helpful discussions for immunohistochemistry, support with microscopy and sharing reagents. We thank the Genomics facility at Institute of Molecular Biotechnology, Austria for providing space and usage of a 10X Chromium controller. Illumina sequencing was done by Ina Nissen, Elodie Vogel Brucklen and Christian Beisel of the Genomics facility at D-BSSE, ETH Zurich. Lentivirus production was done by Oezkan Keles of the Complex Viruses Platform at the Institute of Molecular and Clinical Ophthalmology Basel. FACS sorting support was provided by Mariangela Di Tacchio, Aleksandra Gumienny, Renan Antonialli and Thomas Horn of the Single Cell Facility at D-BSSE, ETH Zurich. Microscopy support was provided by Tom Lummen, Javier Arias Casares and Thomas Horn of the Single Cell Facility at D-BSSE, ETH Zurich. J.G.C., B.T. are supported by CZF2019-002440 from the Chan Zuckerberg Initiative DAF, an advised fund of the Silicon Valley Community Foundation. J.G.C. is supported by the European Research Council (Anthropoid-803441) and the Swiss National Science Foundation (Project Grant-310030_84795). B.T. is supported by the European Research Council (Organomics-758877, Braintime-874606), the Swiss National Science Foundation (Project Grant-310030_192604), and the National Center of Competence in Research Molecular Systems Engineering. J.M.P. has received funding from the T. von Zastrow foundation, the FWF Wittgenstein award (Z 271-B19), the Austrian Academy of Sciences, the Innovative Medicines Initiative 2 Joint Undertaking (JU) under grant agreement No 101005026, and the Canada 150 Research Chairs Program F18-01336, as well as the Canadian Institutes of Health Research COVID-19 grants F20-02343 and F20-02015.

## DATA AND CODE AVAILABILITY

Raw sequencing data will be deposited into ArrayExpress. Processed data and the VCF files for demultiplexing will be deposited in Mendeley Data.

## Methods

### EXPERIMENTAL MODEL AND STUDY DETAILS

#### Pluripotent stem cell lines

In this study we used the human embryonic stem cell (hESC) line H9 obtained from WiCell (Thomson, 1998) and the human induced pluripotent stem cell (hiPSC) line NC8, kindly provided by M. Boehm (Center for Molecular Medicine, National Heart, Lung, and Blood Institute, National Institutes of Health). For CROP-seq experiments, the human iCRISPR 409-B2 cell line, kindly provided by S. Riesenberg and T. Maricic (Department of Evolutionary Genetics, Max-Planck-Institute for Evolutionary Anthropology) was used. This is a doxycycline-inducible Cas9 expressing cell line, generated as described previously (González et al., 2014; Riesenberg and Maricic, 2018).

Karyotyping of all cell lines was carried out using array-based comparative genomic hybridization (aCGH) by the Cytogenetics Laboratory, Cell Guidance Systems Genetics Service. The Agilent ISCA 8×60K v2 array detected no DNA copy number abnormalities in any cell line, except for the NC8 clone used for generation of paired scRNA-seq and scATAC-seq data. This clone showed an approximately 1.7Mb interstitial gain within the proximal long arm of chromosome 20, band q11.21 - a recurrent copy number gain in stem cell lines associated with increased cell survival and proliferation, and an approximately 307kb hemizygous loss within the short arm of chromosome X, band p11.2, which may be constitutional in this sample and not acquired in cell culture. The NC8 clone used for generation of scRNA-seq time course data had intact genomic integrity as shown by multiplex-fluorescence in situ hybridization (M-FISH) and DAPI staining, and by whole-genome single nucleotide polymorphism genotyping array, published previously (Wimmer et al., 2019b).

### METHOD DETAILS

#### Cell culture and human blood vessel organoid (hBVO)generation

##### Pluripotent stem cell culture

All stem cells were cultured under chemically defined, feeder-free conditions and were tested for mycoplasma contamination on a regular basis using EZ-PCR Mycoplasma Detection Kit (Biological Industries, 20-700-20) and found to be negative. Briefly, NC8 and H9 cells were cultured in E8 media (Gibco, A1517001) on 6-well tissue culture plates, pre-coated with 165 µg/ml growth-factor reduced (GFR) Matrigel matrix (Corning, 356231) in DMEM/F12 (Gibco, 11330032), as described previously (Beers et al., 2012). mTagRFPT-CAAX WTC and iCRISPR 409-B2 cells were culture in mTesR Plus (Stem Cell Technologies, 05825) media on 6-well tissue culture plates, pre-coated with hESC-qualified Corning Matrigel matrix (Corning, 354277) in KnockOut-DMEM/F12 (Gibco, 12660012). Media was exchanged daily. Cells were passaged with 0.5 mM EDTA (Invitrogen, AM9261) in phosphate buffered saline (DPBS) (Gibco, 14190250) for regular maintenance. After thawing, cells were passaged at least once before being used for blood vessel organoid generation.

##### hBVO culture

Blood vessel organoids were generated as previously described by Wimmer and colleagues (Wimmer et al., 2019a), with minor alterations. In brief, cells were washed with DPBS (Gibco, 10010023) and subsequently incubated with TrypLE (Thermo Fisher Scientific, 12605010) for 4 min at room temperature. Cells were collected, span down at 200x g for 5 min and resuspended in Aggregation media consisting of KnockOut DMEM/F12 (Gibco, 12660012), 20% KnockOut Serum Replacement (Gibco, 10828028), 1% GlutaMAX (Gibco, 35050061), 1% NEAA (Gibco, 11140035), 55 µM β-mercaptoethanol (Gibco, 21985023), 100 U/ml penicillin-streptomycin, 50 µm Y-27632 (Stem Cell Technologies, 72304). Cells were seeded into Corning Elplasia round-bottom, ultra-low attachment plates at a concentration of 100 to 300 cells per microcavity, depending on the cell line. It has been shown that too small aggregates fail to differentiate or survive, while too large aggregates differentiate inefficiently (Wimmer et al., 2019a). Therefore, we used a microwell plate-format with a defined, standardized number of cells per microwell. Upon the formation of aggregates of size 50 to 100 µm in diameter (in one to two days), media was exchanged for N2B27 media (day 0). N2B27 media consists of Neurobasal (Gibco, 21103049) : DMEM/F12 (Gibco, 11330031) 1:1, 1% B27 supplement (Gibco, 12587010), 0.5% N2 supplement (Gibco, 17502048), 0.5% GlutaMAX (Gibco, 135050061), 55 µM β-mercaptoethanol (Gibco, 21985023), 100 U/ml penicillin-streptomycin (Gibco, 15140122) supplemented with 12 µM CHIR99021 (Tocris Bioscience, 4423) and 30 ng/ml BMP4 (Miltenyi Biotech, 130111165). Three days later (day 3), media was exchanged for N2B27 media supplemented with 2 µm Forskolin (Sigma Aldrich, F3917) and 100 ng/ml VEGF-A (Peprotech, 10020). After two days (day 5), aggregates were embedded in a 12-well plate into Collagen I-Matrigel solution, pH 7.4, consisting of 11.3% v/v 0.1 N NaOH (Sigma Aldrich, S2770), 4.7% v/v 10X DMEM (prepared prior, Sigma Aldrich, D5648-10L), 0.9% v/v HEPES buffer (Gibco, 15630080), 0.7% v/v NaHCO3 (Gibco, 5080094), 0.5% v/v GlutaMAX (Gibco, 25050061), 6.9% v/v Ham’s F12 (Gibco, 11765054), 50% v/v PureCol (Gibco, 15630080), 25% v/v GFR Matrigel (Corning, 356231). From this time on, blood vessel networks and organoids were cultured in blood BVO media - StemPro-34 SFM media, supplemented with StemPro-34 nutrient supplement, 1% GlutaMAX, 100 U/ml penicillin-streptomycin, 15% fetal bovine serum (FBS) (Sigma-Aldrich, F2442-500ML), 100 ng/ml VEGF-A (Peprotech, 10020) and 100 ng/ml FGF2 (Miltenyi Biotech, 1300930841). Media was exchanged fresh every two to three days. Five days after embedding (day 10), gels with blood vessel networks were placed onto a dish lid with sterile lab spoon under a stereomicroscope. Single blood vessel organoids were excised from the matrix manually using sterile 27G needles, and transferred to a 96-well ultra-low attachment plate. BVO media was exchanged fresh every three to four days.

##### hBVO transplantation

Blood vessel organoids were transplanted under the kidney capsule of 12-15-week-old immunodeficient MF1 nu/nu (one H9 and one NC8 organoid per mouse, 8 mice in total) or NOD/SCID/IL2Rγnull (NSG) mice (3 to 6 H9 or NC8 organoids per mouse, 4 mouse each). All animal experiments were performed under ethical animal license protocols from the Austrian Ministry of Science, Research and Economics (BMWFW).

#### Histology

##### Fixation of hBVO

Vascular networks and hBVOs were fixed overnight (12-14 h) at 4°C in 4% weight/volume (wt/vol) paraformaldehyde (PFA) in DPBS. After fixation, samples were washed three times with DPBS at room temperature and stored 4°C until use.

##### Whole-mount immunohistochemistry

Fixed networks and organoids were incubated in Blocking buffer which was DPBS supplemented with 3% volume/volume (v/v) FBS, 1% v/v BSA (Miltenyi Biotec, 130-091-376), 0.5% v/v Triton-X (Sigma-Aldrich, 93420), 0.5% v/v Tween-20 (Sigma-Aldrich, P7949) and 0.01% v/v sodium deoxycholate solution (Sigma-Aldrich, D6750) for 4 h on an orbital shaker at 100 rpm at room temperature. Afterwards, samples were incubated in primary antibodies diluted in Blocking buffer overnight (12-14 h) at 4°C on an orbital shaker at 100 rpm. On the next day, samples were washed three times with PBS-T (DPBS supplemented with 0.05% v/v Tween-20) at room temperature, followed by incubation with secondary antibodies (diluted 1:300) and DAPI (diluted 1:500) in Blocking buffer for 5 h on an orbital shaker at 100 rpm at room temperature. Samples were washed three times with DPBS and mounted in an ibidi glass-bottom chamber (ibidi, 80807) immersed in a drop of ProLong Gold (Thermo Fisher Scientific, P36934). Images were acquired using a spinning disc microscope (Nikon Ti2-Eclipse inverted, Yokogawa CSU-W1 SoRa dual disk dual camera model, T2 spinning disk confocal scanning unit, Yokogawa Uniformizer) with 10X, 20X, 40X silicon oil and 100X immersion oil objectives.

##### Whole-mount multiplexed immunohistochemistry

Whole-mount multiplexed immunohistochemistry was based on the 4i protocol, with slight modifications (Gut et al., 2018). Briefly, blood vessel organoid networks, treated with different pharmacological perturbations, at day 10 of differentiation were fixed with PFA as described above. Tissues were incubated with 400 µL Elution buffer (EB) consisting of 500 mM L-Glycine (Sigma-Aldrich), 5 M Urea (Sigma-Aldrich), 5 M Guanidine chloride (Sigma-Aldrich), 70 mM TCEP-HCl and ddH2O (pH2.5) for 2-4 h on a rocker shaker at room temperature. Afterwards, tissues were rinsed three times with PBS and then washed three times with PBS, for 1 h each, at room temperature. Tissues were then incubated in blocking buffer 1 (BB1) consisting of 5% v/v donkey serum (abcam, ab7475), 1% v/v BSA, 0.5% v/v Triton-X, 0.5% v/v Tween-20 and 0.01% v/v sodium deoxycholate solution, 200 mM NH4Cl (Sigma-Aldrich), 300 mM Maleimide (Sigma-Aldrich) and PBS for 5 h at room temperature. Tissues were then incubated with primary antibodies diluted in blocking buffer 2 (BB2) consisting of 5% v/v donkey serum, 1% v/v BSA, 200 mM NH4Cl, 300 mM Maleimide and PBS for 48 h at 4°C. Tissues were washed three times for 2-4 h with PBS and subsequently incubated with secondary antibodies and DAPI diluted in BB2 for 30 h at 4°C. Samples were rinsed three times with PBS and then washed three times for 2-4 h with PBS at room temperature. All steps described were performed on a rocker shaker.

For imaging, samples were transferred to an 8-well glass-bottom high-wall ibidi chamber and covered with an imaging buffer consisting of 700 mM N-Acetyl-Cysteine (Sigma-Aldrich, A9165-100G) in ddH2O. Images were acquired using a spinning disc microscope, as described above. Afterwards, samples were rinsed once with PBS, then washed three times for 30 min with PBS and again eluted and then stained for a total of two cycles.

##### Antibodies

Primary antibodies used for staining of hBVOs were mouse anti-hCD31 (dilution 1:100, Dako, M0823), rabbit anti-hPDGFR-β (dilution 1:100, Cell Signalling, 3169S), rabbit anti-hAPLNR (dilution 1:100, Invitrogen, 702069), goat anti-hTIE1 (dilution 1:40, R&D, AF619), goat anti-hCollagenIV (dilution 1:150, Millipore, AB769), goat anti-hNRP2 (dilution 1:50, Invitrogen, PA5-47762). Secondary antibodies were donkey anti-mouse AF568 (abcam, ab175472), donkey anti-rabbit AF488 (abcam, ab150073), donkey anti-goat AF647 (abcam, ab150135).

#### Single-cell RNA sequencing (scRNA-seq) of hBVO

##### Cell and tissue dissociation for single cell RNA-seq and single-cell multiome (scMultiome) experiments

Different protocols were used for tissue dissociation, depending on the time point of differentiation. For the generation single cell RNA-seq time course of H9 and NC8 organoid differentiation, a similar number of cells or organoids were pooled together prior to dissociation.

For dissociation of pluripotent stem cell culture, cells were washed once with DPBS and incubated with Accutase (Gibco, A1110501) for 3 to 5 min at 37°C. The reaction was stopped with cold KnockOut DMEM/F12 (Gibco, 12660012) with 10% FBS and span down for 5 min at 300g at 4°C. The cell pellet (on ice) was resuspended in HBSS (without Ca2+, without Mg2+, -/-) with 2% FBS, filtered through a 30 µm cell strainer and centrifuged again for 5 min at 300g at 4°C. The pellet was resuspended in HBSS (-/-) with 2% FBS, counted using Trypan Blue assay on an automated cell counter Countess (Thermo Fisher) and diluted to the appropriate concentration for 10,000 cells to be obtained per lane of the 10x microfluidic chip.

For mesoderm aggregates (age 3 to 5 days), 20-30 aggregates were washed twice with HBSS (without Ca2+, without Mg2+) and incubated in a mixture of one part Accutase and three parts EDTA (Invitrogen, cat. no. AM9261) on a Mini Rocker Shaker at 37°C at 30 rpm for 30 to 40 min, with gentle trituration with a P1000 pipette every 5 min. When most cells were single, the reaction was stopped with cold KnockOut DMEM/F12 (Gibco, 12660012) with 10% FBS and span down for 5 min at 300g in a pre-cooled centrifuge. The cell pellet (on ice) was resuspended in HBSS (without Ca2+, without Mg2+, -/-) with 2% FBS, filtered through a 30 µm cell strainer and centrifuged again for 5 min at 300g at 4°C. The pellet was resuspended in HBSS (-/-) with 2% FBS, counted, diluted to the appropriate concentration if needed and counted again. Cells were loaded on the 10x microfluidic chip aiming for 8,000 cells.

For vascular networks (age 6 to 10 days), one well of a 12-well plate was used. Excess gel was excised with a scalpel, networks were transfer to a 15 ml centrifuge tube, and incubated with cold Cell Recovery Solution (Corning, 35423) for 5 min at 4°C. The networks were triturated gently with a P1000 pipette with a wide-bore tip, centrifuged for 3 min at 300g at 4°C and resuspended in 1 ml pre-warmed enzymatic mix of 5 mg/ml Dispase (Gibco, 17105041) and 0.5 mg/ml Liberase TH Research Grade (Roche, 05401151001) in DPBS (Gibco, 10010023), sterile filtered through a 0.22 µm gaze. The reaction was incubated on a Mini Rocker Shaker at 37°C at 30 rpm for 20 to 30 min. After the first 5 min, and 60 U/ml DNaseI were added and the organoids were triturated gently with a P1000 pipette with a wide-bore tip. Organoids were triturated every 5-7 min and regularly checked under the microscope until a single-cell suspension was achieved. The reaction was stopped with cold KnockOut DMEM/F12 (Gibco, 12660012) with 10% FBS and spun down for 5 min at 300g in a pre-cooled centrifuge. The cell pellet (on ice) was resuspended in HBSS (-/-) with 2% FBS, filtered through a 30 µm cell strainer and centrifuged again for 5 min at 300g at 4°C. The pellet was washed once more with HBSS (-/-) with 2% FBS, then resuspended in HBSS (-/-) with 2% FBS, counted and the concentration was adjusted if necessary. Cells were loaded on the 10x microfluidic chip aiming for 10,000 cells.

For single-cell dissociation of organoids aged 14 days or more, 15 to 20 blood vessel organoids were pooled (20-30 organoids if FACS was to be performed). Organoids were washed once with HBSS (-/-) and subsequently placed onto the lid of a cell culture dish. Excess liquid was aspirated carefully with a 200 µl pipette and organoids were minced with a scalpel. The organoid slurry was transferred to an enzymatic mix of 5 mg/ml Dispase (Gibco, 17105041) and 0.5 mg/ml Liberase TH Research Grade (Roche, 05401151001) in DPBS (Gibco, 10010023). The reaction was incubated on a Mini Rocker Shaker at 37°C at 30 rpm for 20 to 30 min. After the first 5 min, and 60 U/ml DNaseI added and the organoids were triturated gently with a P1000 pipette with a wide-bore tip. Organoids were triturated every 5-7 min and regularly checked under the microscope until a single-cell suspension was achieved. The reaction was stopped with cold KnockOut DMEM/F12 (Gibco, 12660012) with 10% FBS and spun down for 5 min at 300g in a pre-cooled centrifuge. The cell pellet (on ice) was resuspended in HBSS (-/-) with 2% FBS, filtered through a 30 µm or a 40 µm cell strainer and centrifuged again for 5 min at 300g at 4°C. The pellet was washed once more in HBSS (-/-) with 2% FBS, then resuspended in HBSS (-/-) with 2% FBS, counted and the concentration was adjusted if necessary. Cells were loaded on the 10x microfluidic chip aiming for 10,000 cells.

For 10x Multiome experiments, single nuclei were extracted from dissociated vascular networks. After obtaining a single-cell solution, as described above, cells were resuspended in Lysis Buffer containing 10 mM Tris-HCl, pH 7.4 (Sigma-Aldrich, T2194), 10 mM Sodium Chloride (Sigma-Aldrich, 59222C), 3 mM Magnesium Chloride (Sigma-Aldrich, M1028), 0.01% Tween-20 (Bio-Rad, 1662404), 0.01% NP40 Substitute (Sigma-Aldrich, 74385), 0.005% Digitonin (Thermo Fisher, BN2006), 1% BSA (Miltenyi Biotec, 130-091-376), 1 mM DTT (Sigma-Aldrich, 646563), RNase inhibitor (Takara, cat. no. 2313A) and nuclease-free water (Ambion, AM9937). The suspension was incubated for 10 min on ice. The reaction was stopped with Wash Buffer (10 mM Tris-HCl, pH 7.4, 10 mM Sodium Chloride, 3 mM Magnesium Chloride, 1% BSA, 0.1% Tween-20, 1 mM DTT, 1 U/µl RNase inhibitor, and nuclease-free water) and span down for 5 min at 500g at 4°C. The pellet was resuspended in Wash buffer, filtered through a 40 µm Flowmi cell strainer and again span down for 5 min at 500g at 4°C. The pellet was resuspended in 1X Diluted Nuclei Buffer consisting of 1X Nuclei Buffer (stock of 20X, 10x Genomics, 2000207), 1 mM DTT, 1 U/µl RNase inhibitor and nuclease-free water and nuclei concentration was determined. Nuclei were span down for 5 min at 500g at 4°C and resuspended in the respective volume of 1X Diluted Nuclei Buffer according to the 10x Multiome user guide for targeting 10,000 nuclei.

##### scRNA-seq of transplanted hBVOs

For single cell experiments, 8 weeks post-organoid transplantation, mice were sacrificed and organoids were excised from the mouse tissue. 10 to 15 organoids were pooled for each dissociation reaction.

Organoids were collected in a 15-ml centrifugation tube, washed three times with DPBS and transferred to the lid of a 10 cm cell culture dish. Excess liquid was aspirated carefully with a 200 µl pipette and organoids were minced with a scalpel. The organoid slurry was transferred to centrifugation tube with 1 ml pre-warmed enzymatic mix of 30 mg/ml Dispase (Gibco, 17105041) and 1 mg/ml Liberase TH Research Grade (Roche, 05401151001) in DPBS (Gibco, 10010023). The reaction was incubated on a Mini Rocker Shaker at 37°C at 30 rpm for 25 to 35 min. After the first 5 min, and 60 U/ml DNaseI (RNase-free, NEB, M0303L) were added and the organoids were triturated gently with a P1000 pipette with a wide-bore tip. Organoids were triturated every 5-7 min and regularly checked under the microscope until a single-cell suspension was achieved. The reaction was stopped with cold KnockOut DMEM/F12 (Gibco, 12660012) with 10% FBS and spun down for 5 min at 300g in a pre-cooled centrifuge. The cell pellet (on ice) was resuspended in 3 ml Red Blood Cell Lysis Solution (Promega, Z3141) for 4 min and span down for 5 min at 300g in a pre-cooled centrifuge.

For organoids that had been transplanted in MF1 nu/nu mice, the cell pellet was resuspended in HBSS (-/-) with 2% FBS and nuclei were counterstained with DAPI (Sigma-Aldrich, D9542) for 10 min to exclude dead cells during fluorescence-activated cell sorting (FACS). Samples were spun down for 5 min at 300g in a pre-cooled centrifuge and resuspended in HBSS (-/-) with 2% FBS. This step was repeated for a total of two washes. Cells were strained through a 35µm cell strainer and DAPI-negative singlets were sorted on a BD FACS Aria III using BD FACSDiva 8.0.1 Software.

For organoids that had been transplanted in NSG mice, the cell pellet was resuspended in HBSS (-/-) with 2% FBS, and stained against anti-human CD31 conjugated to APC (BD, 558094) for endothelial cells, and anti-human PDGFR-β conjugated to PE (BD, 558821) for mural cells for 45 min. After 35 min, cells were resuspended and nuclei were counterstained with DAPI for additional 10 min to allow dead cell exclusion during cell sorting. Samples were then spun down for 5 min at 300g in a pre-cooled centrifuge and resuspended in HBSS (-/-) with 2% FBS. This step was repeated for a total of two washes. and cells were strained through a 35µm cell strainer. Debri-free single human CD31-or human PDGFR-β-positive cells were sorted on a BD FACS Aria III using BD FACSDiva 8.0.1 Software using a 100 µm nozzle.

##### scRNA-seq data generation

Single-cell experiments were conducted using the 10x Chromium Single Cell 3’ v3 (1000075) or v3.1 (1000121) kit following the manufacturer’s instructions. In brief, cells mixed with reverse transcription mix, gel beads and oil were loaded onto a 10x microfluidic chip to be co-encapsulated into nanodroplets into so-called Gel Beads-in-emulsion (GEMs). Inside each GEM containing a cell, first strand cDNA synthesis occurred, where each mRNA was tagged with a unique molecular identifier (UMI) and a barcode unique for each cell. The droplets were broken, pooled fractions recovered and purified using Dynabeads MyOne Silane (Thermo Fisher, 37002D). cDNA was amplified with a number of PCR cycles depending on the loaded number of cells. Single-cell libraries were generated using fragmentation, end repair, A-tailing and double-sided size selection using SPRIselect Reagent (Beckman Coulter, B23318). P5 and P7 adapters were ligated and individual sample indices provided as a Single Index Plate T Set A (10x Genomics, 2000240) were used to enable pooling and subsequent demultiplexing of multiple libraries. Quantification and quality control of libraries was performed using High Sensitivity DNA assays on an Agilent Bioanalyzer and a Fragment Analyzer. Libraries were sequenced on a NovaSeq 6000 in paired-end mode with 28/8/0/91 cycles.

##### Multiome ATAC-seq and RNA-seq data generation

Single cell Multiome experiments were performed using the 10x Chromium Single Cell Multiome ATAC + Gene Expression (GEX) kit (1000282) following manufacturer’s instructions. Briefly, nuclei suspensions were incubated in a transposition mix where DNA in open regions of chromatin was preferentially fragmented and adapter sequences were added to the ends of the DNA fragments. Afterwards, nuclei were mixed with reverse transcription mix, and gel beads and oil were loaded onto a 10x microfluidic chip to be coencapsulated into nanodroplets, forming GEMs. Inside each GEM containing a nucleus, for GEX, first strand cDNA synthesis occurred, where each mRNA was tagged with a UMI and a barcode unique for each nucleus. In the same partition, for ATAC, transposed DNA was uniquely barcoded and P5 adapters were added. Subsequently, the reaction was quenched. The droplets were broken, pooled fractions recovered and purified using Dynabeads MyOne Silane. Barcoded transposed DNA and barcoded full-length cDNA from polyadenylated mRNA was amplified to fill gaps and generate sufficient amounts for library generation. The pre-amplified product was used for generation of both GEX and ATAC libraries. Single-cell libraries were generated using fragmentation, end repair, A-tailing and double-sided size selection using SPRIselect. P5 and P7 adaptor sequences were ligated and individual sample indices provided as a Dual Index Plate TT Set A (10x Genomics, 3000431) were used to enable pooling and subsequent demultiplexing of multiple libraries. For ATAC libraries construction, P7 adapter sequences and sample indices from a Sample Index Plate N, Set A (10x Genomics, 3000427) were added to the pre-amplified product. For Gene expression libraries, cDNA was further amplified with the number of PCR cycles depending on the number of nuclei loaded. Quantification and quality control of libraries was performed using High Sensitivity DNA assays on an Agilent Bioanalyzer and a Fragment Analyzer. GEX expression libraries were sequenced on a NovaSeq 6000 in paired-end mode with 50/8/24/49 cycles and 28/10/10/90 cycles for GEX and ATAC libraries, respectively.

#### Lineage tracing in hBVOs

##### Generation of iTracer hBVOs and cell scarring

Lineage tracing experiments were performed with the iTracer recording system, kindly provided by T. Gerber and R. Petri, following closely the procedure in the original protocol6. The iTracer plasmid used in this study contained a poly-adenylated and barcoded (11 random bases) dTomato reporter driven by the RPBSA promoter. 91 bases away from the RPBSA promoter, in the opposite direction, is a human U6 (hU6) promoter which drived the expression of gRNA targeting region within the 3’ portion of a dTomato coding sequence.

In brief, iCRISPR 409-B2 cells were cultivated in mTeSR Plus (Stem Cell Technologies, 05825) supplemented with 100 U/ml P/S on Matrigel-coated T75 tissue culture flasks. Cells were electroporated with 10 µg lineage recorder DNA and 1 µg Sleeping Beauty transposase in 100 µl Human Stem Cell nucleofection buffer (Lonza, VVPH-5022) using the B-016 program of the 4D-Nucleofector (Lonza) following the manufacturer’s protocol. Nucleofected cells were plate on Matrigel-coated plates in mTeSR Plus with 100 U/ml P/S and 5 µM Y-27632. Upon reaching around 70% confluency, dTomato-positive cells were sorted on a BD FACS Aria III using BD FACSDiva 8.0.1 Software using a 100 µm nozzle and cultured on Martigel-coated plates in mTeSR Plus supplemented with 100 µg/ml Primocin (InvivoGen, ant-pm-1) and 25 µm Y-27632. After 3 to 4 days, cells were dissociated and used for the generation of blood vessel organoids in one well of a 96-well Elplasia plate, 350 cells per microcavity (27’650 cells in total) as described above.

For scar formation by Cas9, 10 µg/ml doxycycline was added to the blood vessel networks media at day 7. After 24 h media was exchanged with fresh media without doxycycline and organoids were cultured following the protocol described above.

##### scRNA-seq of iTracer hBVOs and barcode and scar library preparation

At day 14 of differentiation, organoids were dissociated, processed for generation of scRNA-seq libraries and sequenced on a NovaSeq 6000, as described above.

Additional enrichment polymerase chain reactions (PCRs) were performed where barcode and scar regions from 60 ng of cDNA remaining from the scRNA-seq preparation were amplified in three separate reactions. First, cDNA was amplified in a PCR broadly targeting the barcode and scar region. Subsequently, two 10 ng portions of the reaction product were used for further separate reactions, one for scar and one for barcode amplification. Finally, Illumina adapter sequences were added. After every PCR, samples were purified with SPRIselect beads (Beckman Coulter, B23319). The libraries were sequenced on a NovaSeq 6000. Primers for the PCRs can be found in Table S9.

#### CROP-seq in hBVOs

##### Library design, vector and lentivirus preparation for perturbation experiment

The perturbation experiment was based on the CROP-seq protocol (Datlinger et al., 2017) with minor alterations. The CROP-seq-EGFP vector – a modified version of the original CROP-seq vector where the Puro selection marker was exchanged with an EGFP selection marker, was kindly provided by J. S. Fleck, D. Wollny and M. Seimiya. Therefore, iCRISPR 409-B2 cells, which contain a selection marker, could be efficiently transfected with the vector and selected for the expression of EGFP via FACS.

18 TFs and 16 receptors were chosen as target genes. Two libraries were generated, each targeting 17 different genes and containing 8 non-targeting gRNAs. Three gRNAs per target gene were designed using the online tool CRISPick (Broad Institute, https://portals.broadinstitute.org/gppx/crispick/public) and synthesized by IDT as 74 base oligonucleotides with 19 and 35 bases of homology to the hU6 promoter and guide RNA backbone, respectively. Oligonucleotides were diluted to 100 µM and pooled in equal amounts for each library. gRNA libraries were cloned by Gibson’s isothermal assembly. gRNA complexity validation was performed by attaching Illumina i5 and i7 indices to 10 ng of the pooled plasmid library via PCR and sequencing on an Illumina MiSeq platform.

Upon validation of the libraries, lentiviral packaging was performed by the Viral Core Facilty at the Institute of Molecular and Clinical Ophthalmology Basel.

##### Evaluation of genome editing

To verify and quantify the efficiency of CRISPR/Cas9-mediated knock-out, we screened for insertions and deletions (INDELs) in the cell population using a mismatch cleavage assay that utilizes the T7 endonuclease I (T7EI). We performed the assay for gRNAs targeting MECOM, ETV2 and PITX1 (three guides per gene).

In brief, 3 µL of a stock of 100 µM Alt-R CRISPR-Cas9 crRNA (IDT) and 3 µL of a stock of 100 µM Alt-R CRISPR-Cas9 tracrRNA (IDT, 1072533) were diluted with 94 µL Nuclease Free Duplex Buffer (IDT, 11010301). The crRNA:tracrRNA mix was incubated at 95°C for 5 min in a thermocycler after which they were allowed to cool down at room temperature.

Lipofectamine transfection complex was prepared by mixing 0.75 µL of 3 µM gRNA, 0.38 µL of Lipofectamine RNAiMAX (Thermo Fisher Scientific, 13778100) and 23.87 µL of OptiMEM media (Thermo Fisher Scientific, 51985091) per reaction. The complex was incubated for 20 min at room temperature. In the meantime 409-B2 iCRISPRn (iCas9 cells) which had been incubated for 24 h with 2 µg/mL Doxycycline, were dissociated with the following procedure: cells were washed once with DPBS, incubated with TrypLE (Thermo Fisher Scientific, 12605010) for 3 min at 37°C. TrypLE was then aspirated, cells were collected with 2 mL mTeSRplus (Stem Cell Technologies, 05825), span down and resuspended in 1 mL mTeSRplus. Cells were counted and per well of a 96-well plate, 40,000 cells were seeded in 150 µL media consisting of 2 µg/mL Doxycycline in 25 µL Lipofectamine complex, 15 µL CloneR (Stem Cell Technologies, 5889), and 85 µL mTeSRplus.

Media (mTeSRplus supplemented with 100 U/mL P/S) was exchanged daily. When cells reached a confluency of 80%, media was aspirated, cells were washed once with DPBS and genomic DNA (gDNA) was extracted with 50 µL/well of Quick-Extract solution (Lucigen, QE09050). Lysates were pipetted up and down thoroughly, incubated in PCR tubes at 65°C for 6 min, followed by 98°C for 2 min, then frozen and stored at -20°C until use.

For the T7EI assay the gDNA of the edited cell population was used as a direct PCR template for amplification with primers specific to the target region (done for each gRNA separately; see Table S6). The PCR reaction consisted of 10 µL Phusion High-Fidelity PCR MasterMix 2x (Thermo Fisher Scientific, F531S), 1 µL of the forward and reverse primer, each, 1 µL of the gDNA template, and 7 µL nuclease-free H2O. The PCR program consisted of: 30 s at 98°C, 35 cycles of (10 sec at 98°C, 15 sec at 64°C, 10 sec at 72°C), 2 min at 72°C in a BioRad C1000 Thermal Cycler. The PCR product was cleaned with SPRI beads (Beckman Coulter, B23319) size selection (0.9x) and eluted in 30 µL Elution Buffer. To produce heteroduplex mismatches where double-strand breaks have occurred, 50 ng PCR product was incubated with 2 µL 10X NEBuffer 2 (NEB, B7002S) and nuclease-free water adding up to 19 µL using the following program: 5 min at 95°C, at 95-85 with ramp speed of -2°C/s, at 85-25°C with ramp speed of -0.1°C/s in a BioRad C1000 Thermal Cycler. These mismatches are then recognized and cleaved by T7EI in a heteroduplex digestion, where 19 µL of the annealed PCR product were incubated with EnGen T7 Endonuclease I at 37°C for 15 min, followed by the addition of 1 µL Proteinase K, and further incubated at 37°C for 5 min to inactivate the T7 enzyme.

Fragment analysis was performed with a High Sensitivity DNA kit on a Bioanalyzer following the manufacturer’s instructions.

##### Generation of mosaic hBVOs with genetic perturbations

For generation of mosaic perturbed vascular organoids, 409-B2 iCRISPR cells were grown on Matrigel-coated plates in mTeSR Plus (Stem Cell Technologies, 05825) and dissociated into single cells using TrypLE (Thermo Fisher Scientific, 12605010) when at 60% - 80% confluency. 2.1 Mln cells in mTeSR Plus supplemented with 100 U/ml P/S and 5 µm Y-27632 were plated per T75 tissue culture flask pre-coated with Matrigel. After 24 h, cells were transduced at a low multiplicity of infection (MOI), so that no more than 3% of the cells were GFP-positive in order to reduce the possibility of cells receiving more than one lentivirus (and therefore, one gRNA). The virus was added to the cell culture media consisting of mTeSR Plus, 100 U/ml P/S and 5 µm Y-27632. 24 h later media was exchanged to mTeSR Plus and 100 U/ml P/S. After 2 to 4 days, GFP-positive cells were sorted on a BD FACS Aria III using BD FACSDiva 8.0.1 Software using a 100 µm nozzle and subsequently cultured on Martigel-coated plates in mTeSR Plus supplemented with 100 µg/ml Primocin and 25 µm Y-27632. 24 h later media was exchanged to mTeSR Plus supplemented with 100 µg/ml Primocin, 5 µm Y-27632 and 2 µg/ml doxycycline (Sigma-Aldrich, D9891). For the next 4 to 6 days, until reaching 70% confluency, cells were cultured in mTeSR Plus supplemented with 100 µg/ml Primocin and 2 µg/ml doxycycline. Blood vessel organoids were generated either from GFP-positive cells only (samples in batches 1 and 2) or from 20% GFP-positive cells and 80% non-transduced cells (samples in batch 2).

##### scRNA-seq of mosaic hBVOs with genetic perturbations

scRNA-seq experiments were performed with organoids at day 7, 14, or 21 following the procedure described in the dissociation into single cells subsection and the scRNAseq GEX libraries generation section of this study. Two batches of organoids with multiple samples per batch were run. Organoids from batch 1 were dissociated and cells were directly loaded onto a 10x Genomics chip for subsequent scRNA-seq. Organoids from batch 2 were dissociated, GFP-positive cells were sorted on a BD FACS Aria III using BD FACSDiva 8.0.1 Software using a 100 µm nozzland and only GFP-positive cells were loaded onto a 10x Genomics chip.

##### Enrichment of gRNA-containing transcripts from scRNA-seq

gRNA was amplified from 60 ng cDNA remaining from the scRNA-seq preparation in three separate PCRs, similar to others (Fleck et al., 2021; Hill et al., 2018). First, the gRNA region was amplified in a PCR broadly targeting the outer part of the human U6 promoter. Second, the inner part of the U6 promoter was amplified and a Truseq Illumina i5 adapter was added. A Truseq Illumina i7 adapter was added in the last reaction. After every PCR, samples were purified with SPRIselect beads (Beckman Coulter, B23319). The libraries were sequenced on a NovaSeq 6000. Primers for the PCRs can be found in Table S5.

#### Signaling pathways modulation of blood vessel organoids

##### Generation of hBVOs with pharmacological perturbations

For studying the importance of Notch and VEGF signaling pathways in blood vessel development, we performed a screen with 22 different conditions (see Table S7 for details), varying the concentration of Forskolin and VEGF and testing the effect of DAPT, a γ-secretase inhibitor (Sigma-Aldrich, D5942) on blood vessel organoid development.

##### RNA extraction from pharmacologically perturbed hBVOs

Blood vessel organoid networks treated with the different conditions and 2 control samples (original protocol) were enzymatically dissociated at day 14. Two individual biological replicates of each condition and control were cultured and treated separately for initial test and in addition one more biological replicate was generated for testing conditions which were of interest based on the first PCR. Each well was incubated with pre-warmed enzymatic mix of 6 mg/ml Dispase (Gibco, 17105041) and 1 mg/ml Liberase TH Research Grade (Roche, 05401151001) in DPBS (Gibco, 10010023) for 8 min at 37°C. 8 U/ml DNaseI (RNase-free, NEB, M0303L) were added to each sample, followed by gentle trituration with a wide-bore P1000 pipette and further incubation for 6-7 min at 37°C. The reaction was stopped by adding cold KnockOut DMEM/F12 (Gibco, 12660012) with 10% FBS and centrifugation for 2 min at 300g in a pre-cooled centrifuge (4°C). Samples were washed once with DPBS, and spun down for 2 min at 300g in a pre-cooled centrifuge. The supernatant was discarded, cold TRIzol (Invitrogen, 15596026) was added to each pellet (200 µl per 24-well) for RNA extraction and samples were triturated gently 8-10 times. Extracts were used immediately or stored at -80°C. Further RNA purification was performed using Direct-zol RNA Miniprep (ZymoResearch, R2050) according to manufacturer’s instructions.

The RNA concentration and purity were evaluated with a NanoDrop 2000 spectrometer (Thermo Fisher Scientific).

##### RT-qPCR of pharmacologically perturbed hBVOs

50 ng RNA of each sample was reversely transcribed into cDNA using ReadyScript cDNA Synthesis Mix (Sigma-Aldrich, RDRT), with the following cycler programme: 5 min at 25°C, 30 min at 42°C, 5 min at 85°C, hold at 4°C in a C1000 Thermal Cycler (Biorad). We then performed RT-qPCR for gene expression quantification of the following genes: CXCR3, EFNB2, GJA4, EPHB4, APLNR, NR2F2, PECAM1, PDGRFB, ACTA2, GAPDH (used as an internal normalization control) (see Table S7 for primer sequences, ordered from IDT). RT-qPCR was performed using KAPA SYBR FAST qPCR Kit (Kapa Biosystems, KK4602) according to manufacturer’s instructions with the following cycler programme: 3 min at 95°C, 45 cycles of (3 sec at 95°C, 20 sec at 60°C, 10 sec at 72°C), 10 sec at 95°C, 1 min at 65°C, 1 sec at 97°C in a LightCycler96 (Roche).

##### scRNA-seq of pharmacologically perturbed hBVOs

For scRNA-seq experiments, 10-day-old blood vessel organoid networks were dissociated into single cells as described in the tissue dissociation for scRNA-seq section of this article and loaded on the 10x microfluidic chip aiming for 10,000 cells.

### QUANTIFICATION AND STATISTICAL ANALYSIS

#### scRNA-seq data processing and analysis

##### Time course hBVO scRNA-seq data preprocessing and analysis

Cell Ranger (10x Genomic, v4.0.0) was used to map the sequencing reads to the human reference (GRCh38-based, 10x Genomics, v3.0.0) and call cells to generate the count matrices. Demultiplexing of lines was done with demuxlet (Kang et al., 2018), given the genotypes of H9 and NC8 lines in VCF files and the BAM files produced by the mapping. The count matrices of exonic/intronic reads were produced with dropEst (Petukhov et al., 2018), given the same gene annotation as the CellRanger reference.

Next, the scRNA-seq data representing different lines and time points were merged and normalized using Seurat (v. 4.0.0) (Hao et al., 2021). Additional quality control was done by filtering cells with fewer than 500 genes detected or with mitochondrial transcript percentage higher than 15%. The top-5000 highly variable genes were defined using the vst method (Table S2). Among them, mitochondrial genes, ribosomal genes and cell-cycle-related genes were excluded. The expression values of each highly variable gene across cells were scaled, after regressing out the cell-cycle scores (G2M and S) calculated by Seurat. Principal component analysis (PCA) was applied to the scaled expression matrix for the first 20 principal components (PC). Cluster similarity spectrum (CSS) (He et al., 2020) was applied to integrate samples, i.e. lines+timepoints. PCA was applied to the resulting CSS matrix to further compress the integrated space to 20 dimensions, which were then used to generate the Uniform Manifold Approximation and Projection (UMAP) embedding, as well as clusters using louvain clustering (resolution = 0.8). Connectivities between clusters were quantified and tested based on the k-nearest-neighbor (kNN, k = 20) graph of cells calculated based on the PCA-reduced CSS space. In brief, the number of edges in the cell kNN graph between cluster *i* (with cell number *n*_*i*_) to cluster *j* (with cell number n_j_) was counted as N_ij_. The connectivity from cluster *i* to cluster *j* was then tested by comparing *n*_*j*_*/N*_*ij*_ and 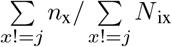 using Fisher’s exact test. Annotations of the resulting clusters were done based on the combinatorial expression of canonical cell type markers and *de novo* cluster markers. The cluster markers were identified using the presto package by combining multiple criteria (Benjamini-Hochberg (BH)-adjusted Wilcoxon test P < 0.01, logFC > 1.2, AUC > 0.7, detection rate difference > 20%). RNA velocity analysis was done with scVelo (Bergen et al., 2020), using the PCA-reduced CSS space to represent transcriptome of cells.

To investigate gene expression changes along the endothelial cell (EC) differentiation trajectory, related cells which include clusters annotated as lateral plate mesoderm (c4, c13, c9, c10, c2), EC (c14, c15, c11, c18, c5, c6, c20), and one mural cell (MC) cluster (c7) were subsetted. Louvain clustering (resolution = 2) was applied to the subsetted cells to identify one outlier cluster (subcluster 26), which was excluded from the following analysis. Diffusion map pseudotimes (DPT) of the remaining cells were obtained using the destiny R package (Angerer et al., 2016), which was applied to the PCA-reduced CSS space directly. The resulting DPT were ranked to retrieve the final EC differentiation pseudotime. To identify genes with expression changed along the reconstructed pseudotime series, an F-test-based ANCOVA-like test was applied as described previously (Kanton et al., 2019). In brief, for each gene detected in more than 1% of cells, two linear models were fitted. The full model includes the natural splines of the pseudotime (degree of freedom (d.f.) = 5) and the line information as the covariate; while the reduced model includes only the line information. A one-sided F-test was then applied to compare the mean squared errors of the two models. Genes with Bonferroni-corrected P < 0.05 were considered with pseudotime-dependent expression along the EC differentiation trajectory. Hierarchical clustering was applied to the correlation distance of the resulting genes across their average expression in each of the 20 pseudotime bins, each of which contained 5% of the cells (Table S2).

##### scMultiome data preprocessing and analysis

Cell Ranger ARC (10x Genomics, v2.0.0) was used to map the sequencing reads of both the RNA and ATAC libraries to the human reference (GRCh38-based, 10x Genomics, v2020-A-2.0.0) and call cells. The data of two lines (H9, NC8) with both modalities (RNA/ATAC) were merged with Seurat (v4.0.0) and Signac (v1.4.0) in R. For the RNA modality, SCTransform (Hafemeister and Satija, 2019) was used to normalize the data, followed by PCA to obtain 20 PCs. CSS was applied to integrate the two lines, followed by PCA on the CSS space to obtain the PCA-reduced CSS space (20 dimensions). UMAP embedding was generated given the PCA-reduced CSS space as the input using default parameters. For the ATAC modality, peaks were firstly re-called using Signac by calling MACS2 (v2.2.7.1) by combining data of both cell lines, and then requantified in each cell with Signac. The ATAC data were then normalized as TFIDF using Signac (default parameters) and partial singular value decomposition (SVD) was applied to the normalized values of peaks with fragments detected in at least five cells to obtain the latent semantic indexing (LSI) representation of the data. The 1^st^ LSI was discarded as it is highly correlated with the ATAC library coverage of cells. Similar to the RNA modality, CSS was applied to integrate the ATAC modality of the two lines, with UMAP embedding generated given the PCA-reduced CSS space of the ATAC data using the default parameters. To jointly consider both RNA and ATAC modalities, the weighted nearest neighbor (WNN) graph was constructed using Seurat, given the PCA-reduced CSS spaces of the two modalities as the input. The resulting WNN graph was used to generate the bi-modal UMAP embedding (default parameters), as well as to do the Louvain clustering (resolution = 0.05) to get the three main clusters, which were annotated as EC, MC, and others by considering expression of the canonical markers.

To project the scMultiome data to the time course transcriptomic atlas, the CSS model representing the time course data was applied to the RNA modality of the scMultiome data, resulting in its projected CSS matrix. The projected UMAP embedding of the scMultiome data to the time course atlas was obtained by using the ProjectUMAP function in Seurat, using the CSS representation of the time course data and the project CSS representation of the scMultiome data as the input. The time course cluster labels were transferred to cells in the scMultiome data using a kNN (k = 50) classifier (He et al., 2020).

To identify peaks showing significant accessibility differences among different time course cell clusters, an ANCOVA-like method based on the generalized linear model (GLM), similar to described before (Kanton et al., 2019), was used. In brief, two GLMs with binomial error distribution and a logistic link function were fitted for each peak detected in more than 0.5% of cells. The full model includes the projected cluster labels and the line information as covariate, while the reduced model only contains the line information. The two models were compared using Chi-square test. Peaks with Bonferroni-corrected P < 0.01 were considered with cluster-variable accessibility, namely cluster-variable peaks. For each of the cluster-variable peaks, its model coefficients of each cluster in the full GLM model were seen as its relative accessibility in each cluster. The accessibilities of different clusters were then ranked for each cluster-variable peak, and a kNN (k = 20) graph of cluster-variable peaks was reconstructed based on the Euclidean distances between the ranked accessibility profiles of peaks. Co-accessibile peak modules were obtained by applying Louvain clustering (resolution = 0.5) to the cluster-variable peak kNN graph. The GREAT functional enrichment analysis was applied to each co-accessible peak module, given all the called peaks located at the assembled chromosomes as the background.

To reconstruct the gene regulatory network (GRN) governing the cell type identity maintenance in BVO, we applied Pando (Fleck et al., 2021) to the scMultiome data, using the default parameters. In brief, Pando firstly incorporates the peak coordinates, evolutionary property of genomic regions, pre-defined cis-regulatory elements, transcription factor (TF) motif databases and prediction to identify putative TF binding sites regulating each gene. Afterwards, a linear model was fitted for each gene on its (log-normalized) expression based on all candidate TF-binding site interactions. The fitted coefficients were tested for significance using ANOVA, followed by BH multiple test correction to obtain an FDR-adjusted P value, to which a significance threshold of 0.05 was applied.

To visualize the reconstructed GRN of TFs, the pairwise Pearson correlation coefficients (PCCs) between TFs were firstly calculated as the base TF-TF linkage score. Next, the lineage score between any TF pair with no inferred direct regulatory relationship was set to 0. A partial PCA was then applied to convert the linkage score matrix to represent each TF by a 20-dimensional vector. This truncated PC matrix was used as the input to generate the UMAP embedding of the TFs.

##### Preprocessing and analysis of the hBVO iTracer data

For both the iTracer and scRNA-seq libraries, we used Cell Ranger (10x Genomics, v4.0.0) to align them to the modified human reference (GRCh38-based, 10x Genomics, v3.0.0) with the RFP sequence added using the default parameters. The iTracer barcode and scar information was retrieved as described before (He et al.). In brief, the iTracer barcode transcripts were firstly filtered, excluding UMIs with no more than three reads. A loess model of number of reads per UMI against the frequency (loess (nreads ≈ log (freq_nreads))) was then fitted. UMIs with fewer reads than the first minimum of the fitted model were discarded. Additional requirements, including one barcode per UMI, barcodes beginning and ending with A/T, only one barcode-UMI being allowed in one cell, were also applied. Similar quality control pipeline was also applied to the iTracer scar transcripts. Lastly, any barcode or scar transcript without any corresponding transcript in the other library were excluded. Based on the iTracer readout, we defined a barcode family as the union of cells with the same barcode combination detected. Cells in the same barcode family were likely expanded from the same iPSC. In each barcode family, scar families, that is, cells with the same scar combination, were further defined, which represents cells in the organoid expanded from the same cell when Cas9 was induced.

For the transcriptomic data, additional quality control was performed by excluding cells with no more than 1500 genes detected or mitochondrial transcript percentage larger than 10%. Seurat (v4.0.0) was used to perform normalization, highly variable gene identification (vst method, 2000 genes; Table S2) and scaling, PCA for dimension reduction, UMAP embedding reconstruction, as well as Louvain clustering (resolution = 0.5). Clusters were annotated as EC or MC based on the marker expression. The cell type annotation and lineage information inferred from the iTracer readout were incorporated to look for clones generated from the bi-potent stem cells at day 7 of the BVO culture.

##### Preprocessing and analysis of the hBVO CROP-seq data

Similar to described above, Cell Ranger (10x Genomics, v4.0.0) was used to generate transcriptomic count matrix of the hBVO CROP-seq data. Separated quality control based on detected gene number and mitochondrial transcript percentage was firstly applied to different samples from the two experimental batches. The scRNA-seq data of these samples were then merged, followed by normalization, highly variable gene identification (vst method, 3000, cell-cycle-related genes, mitochondrial genes and ribosomal genes excluded; Table S2), data scaling (cell-cycle scores regressed out) and PCA. CSS was further applied to integrate data of different samples, followed by PCA on CSS to obtain the PCA-reduced CSS representation (with 10 dimensions), which was then used to generate the UMAP embedding as well as Louvain clustering (resolution = 2). The resulting clusters were annotated as EC, MC, hematopoietic cells, as well as other cell types such as neural crest derivatives, surface ectoderm and neuromesodermal cells, based on the combinatorial expression of canonical cell type markers.

To assign gRNA labels to cells, reads obtained from the amplicon sequencing were firstly mapped to the extended human reference (GRCh38-based, 10x Genomics, v3.1.0) using Cell Ranger (v4.0.0) with the artificial chromosomes representing the Guide-GFP construct used in the CROP-seq experiment (Datlinger et al., 2017). Similar quality control procedure as described before (Fleck et al.) was used to extract informative gRNA transcripts detected in each cell based on a Gaussian mixture model of number of reads per UMI, resulting in a cell x guide count matrix, which was further binarized to obtain the final cell-to-gRNA assignments.

We used a Cochran-Mantel-Haenzel (CMH) test stratified by sample to assess the effect of gene knock-out (KO) on cell type composition in hBVO. This allows the control of confounding effects through differential gRNA abundance in different samples. The test was applied to each gRNA targeting different genes separately using the pool of dummy gRNAs, comparing their frequency in one cell type relative to the others. The test returns the estimated odds ratio (> 1 for enrichment, < 1 for depletion), as well as the P value for statistical significance. BH correction was applied to the P values to correct for the multiple testing. To assess the robustness of the identified effect in the two batches, the test was also applied to samples from each of the two batches separately. In order to derive the gene-level estimates, we also applied the same test to the pool of different gRNAs targeting the same gene.

To assess the compositional changes induced by the target gene KO in a finer resolution and cluster-free manner, the kNN (k = 100) graph of cells was firstly constructed based on Euclidean distance on the PCA-reduced CSS space. Next, a CMH test stratified by batch was performed on the neighborhood of each cell, comparing frequencies of the gRNA or gRNA pool and the pool of dummy gRNAs within and outside of the neighborhood. The resulting neighborhood enrichment score of each cell was defined as signed -log(P), where the sign was determined by the sign of log-transformed odds ratio. A random walk with restart procedure (*s*^*t*^*=*β *x s* ^*0*^*+(1-*β*)Ws*^*t-1*^, β*=0*.*1, t* ≤ *100*) was then applied to smooth the neighborhood enrichment score of each cell. Here,*s*^*0*^ was the vector of original neighborhood enrichment scores of all cells, and *W* is the transition matrix defined as the column-normalized adjacency matrix of the kNN graph. Visualization of the results was done using schex (Freytag and Lister, 2020) which implements hexagonal binning for single-cell data visualization.

To assess the target gene KO effect on transcriptomic states of cells, an ANOVA-based method was used. In brief, a linear model was fitted for each gene detected in more than 1% of the cells. The full model was fitted on the log-transformed gene expression values against perturbation status and sample information as the covariate. ANOVA was then used to assess the significance of variation explained by the perturbation status. The test was applied to each target gene in EC and MC separately. Only cells with detected gRNAs targeting the same gene, as well as those with only dummy gRNAs were considered in the analysis. Here, the perturbation status was represented by the inferred perturbation probability as described (Fleck et al., 2021).

##### Preprocessing and analysis of the transplanted hBVO scRNA-seq data

Cell Ranger (10x Genomics, v4.0.0) was used to align reads of the scRNA-seq data to the human-mouse dual genome reference (GRCh38 and mm10, 10x Genomics, v3.1.0) to demultiplex cells from different species. The mapping to the human reference (GRCh38, 10x Genomics, v3.1.0) was then performed, and the count matrix for each sample was generated with only cells assigned as human cells in the species demultiplexing included. Cells with fewer than 1500 genes detected, more than 6000 genes detected, as well as those with mitochondrial transcript percentage >15%, were also filtered. Seurat was then used to perform normalization, highly variable gene identification (vst method for 3000 genes, followed by excluding cell-cycle-related, mitochondrial, ribosomal genes; Table S2), data scaling with cell-cycle scores regressed out, and PCA. CSS was used to integrate data of different samples (cluster_resolution = 1), followed by further dimension reduced by truncated PCA (top-20 PCs). The PCA-reduced CSS representation was used to generate the UMAP embedding (default parameters) and perform Louvain clustering (resolution = 0.8). Annotation of the clusters into EC or mesenchymal cell (MC) categories was done based on the combinatorial expression of canonical cell type markers. The data was further split into the EC (cluster 6 and 9) and MC (other clusters excluding cluster 18) portions for further analysis. Similar pipeline as described above was then applied to the EC-only portion again for further characterization, with the UMAP embedding remade and Louvain clustering (resolution = 0.2) redone on the recalculated PCA-reduced CSS representation. Cluster markers were identified using the presto package, requiring multiple criteria including FC > 1.2, AUC > 0.65, BH-corrected P < 0.01, detection rate difference > 20

##### Transcriptome-based GRN reconstruction using SCENIC

We used SCENIC (Aibar et al., 2017) to infer the GRN controlling the transcriptomic changes during BVO development and upon BVO transplantation. In brief, we considered all genes detected in more than 1% of cells merging the time course hBVO scRNA-seq data and the transplanted BVO scRNA-seq data. Next, 10,000 cells were randomly subsetted from the merged data, and pySCENIC (v0.11.2) was applied to the subset with grn-boost2 method and other default parameters to obtain the regulome of TFs. It is worth mentioning that only positive regulomes are inferred with the used default parameter sets.

The AUCell score of each inferred regulome in each cell was also calculated by pySCENIC for the cell subset used to infer the regulomes, and was used as the proxy of the regulome activity. The AUCell scores of the other cells were then inferred based on the calculated ones using an iterative network diffusion method. In brief, a kNN graph was reconstructed for the time course and transplanted hBVO scRNA-seq separately. For each of the two data sets, the row-normalized (sum of each row is 1) adjacency matrix of the kNN graph was modified as the diffusion transition matrix (*W*), where *W*_*i,i*_*=1* and *W*_*i,j*_*=0 (j*≠*i)* if and only if cell i is among the training subset. The iterative diffusion scheme was then performed as *S*^*t*^*=WS*^*t-1*^ *(t* ≤ *30)*, where *S*^*0*^ is the initial cell x regulome AUCell matrix with the uncalculated AUCell scores set to 0. Afterwards, another iterative diffusion scheme (5 rounds) was applied to further smoothen the estimated cellular regulome activities, with the row-normalized unmodified kNN graph adjacency matrix used as the diffusion transition matrix.

##### Quantification of the signaling pathway activity

We retrieved the gene list annotated in each KEGG pathway of humans via KEGG API using the R package KEGGREST. Pathways with names matching the pattern “signaling pathway” were kept as the signaling pathways.

To quantify the activity of each pathway in a cell, we developed the following procedure to summarize expression of genes involved in the pathway. Firstly, any ligand genes according to the CellPhoneDB database (v2) were excluded. Next, for any TF in the pathway which has the regulome inferred in the SCENIC analysis, the AUCell score of the TF regulome was used as the proxy of the downstream effect of the pathway. The signature matrix of the pathway in all cells was then generated by concatenating the scaled expression matrix of the non-ligand genes in the pathway (cell x gene) and the scaled regulome activity matrix of TFs in the pathway (cell x regulome). PCA was applied to the signature matrix, and the first PC (PC1) was seen as the summary score. Lastly, the summary score was correlated with genes and regulomes in the signature matrix. If more than half of the individual signatures show negative correlation with the summary score, the sign of the summary score was flipped. The resulting summary score was used as the metric of signaling pathway activity.

The signaling pathway activity was firstly trained and calculated on the time course and transplanted BVO scRNA-seq merged data. The parameters in the model to calculate the summary score of each signaling pathway, including mean and standard deviation of each signature for data scaling as well as the PC1 loading vector, were saved so that the same model can be applied to other data to generate comparable summary scores.

##### Preprocessing and analysis of the signaling pathways modulation screen scRNA-seq data

Similar procedure as described above was used to preprocess the scRNA-seq data of the signaling pathways modulation screen. During the quality control of cells, those with < 1000 genes detected or mitochondrial transcript percentage > 25% were filtered. Cell cycle scores and phase inference was done with Seurat. 5000 highly variable genes were defined (Table S2). UMAP embedding was generated and Louvain clustering (resolution = 0.8) was performed based on the first 30 PCs in PCA. The resulting clusters were grouped into EC and MC based on the canonical cell type marker expression.

Next, cells in the EC clusters were subset for further analysis. 3000 highly variable genes were redefined (Table S2), followed by excluding cell-cycle related, mitochondrial and ribosomal genes. Data scaling was done after regressing out the cell cycle scores, and 20 PCs were calculated based on the scaled expression of the highly variable genes. CSS was then used to integrate data of different conditions, followed by truncated PCA to further reduce the number of dimensions to 20. A linear model of each reduced dimension was then fitted against the cell cycle scores. The residuals of the 20 models were concatenated to form the final cell-cycle-regressed PCA-reduced CSS representation, which was then used to generate the UMAP embedding and perform Louvain clustering (resolution = 0.2). As one cluster (c2) showed PDGFRB^+^ which implies the possibility of being doublet, cells in that cluster were filtered, and the same pipeline was applied again to the remaining cells. The final clustering (resolution=0.5) was performed on the re-calculated cell-cycle-regressed PCA-reduced CSS representation. Cluster markers were identified using the presto package, requiring FC > 1.2, AUC > 0.6, BH-corrected Wilcoxon test P < 0.01, detection rate difference > 20% and detection rate outside of the cluster < 30%. AUCell scores of the SCENIC-inferred regulomes were calculated for the data set, and the signaling pathway activity scores were also calculated using the models obtained as described above.

Similar analysis was also applied to the MC subset of the data to characterize heterogeneity differences of MC in different signaling pathway modulation conditions.

##### Comparison to the public data sets

Two public data sets were used in this study. The scRNA-seq atlas of the developing human gut tube (Yu et al., 2021) was retrieved from Mendeley Data (https://doi.org/10.17632/x53tts3zfr.1), which includes the major cell type annotation as well as clusters, and was directly used to benchmark the major cell type identity of the time course and transplanted BVO scRNA-seq data. In addition, the mesenchymal cell subset of the same atlas, which includes detailed mesenchymal subtype annotations, was obtained from the same site and used to help annotate the mesenchymal cell clusters in the transplanted BVO scRNA-seq data.

Moreover, we also subsetted endothelial cells from the full atlas for further characterization. Using Seurat, 3000 highly variable features were firstly identified (vst method; Table S2), with data scaling applied to those genes. PCA was performed (20 PCs calculated). CSS was used to integrate cells from different samples, followed by PCA for further dimension reduction (10 PCs). UMAP embedding was generated based on the PCA-reduced CSS representation, so as the Louvain clustering (resolution=0.5). Annotation of clusters were done by incorporating tissue origins of cells and the combinatorial expression of cell type markers. Arterial and venous EC signature genes were generated by differential expression analysis between the “Arterial EC” and “Venous EC” clusters using the presto package, requiring FC > 1.2, AUC > 0.7, BH-corrected Wilcoxon test P < 0.01, detection rate difference > 20% and detection rate outside of the cluster < 20%. Those genes were used to calculate the arterial and venous signature scores, respectively, using the AddModuleScores function in Seurat. The AV specification score was calculated as the arterial signature score subtracted by the venous signature score.

To study the organ specificity of ECs, the EC subset of the human cell atlas of fetal gene expression (Cao et al., 2020) was retrieved. The data was then split based on the organ origins of cells. For each organ subset, similar procedures as follows were applied. Using Seurat (v4.0.0), data normalization was performed, and 2000 highly variable genes were identified (vst method). DAVID enrichment analysis was performed on those highly variable genes, given all genes detected in the data as the background. Data scaling was done after regressing out the cell-cycle scores calculated by Seurat. PCA was performed, and using the first 20 PCs Louvain clustering with varied resolution was performed. The resulting clusters were then annotated based on expression of canonical EC subtype markers, or represented by their top markers if the canonical markers did not show a clear pattern. Transcriptomic similarity between ECs in the *in vitro* or transplanted BVO and ECs from different organs was calculated as the Pearson correlation coefficient between average expressions of the 2000 highly variable genes in different EC subtypes in different organs, and those in the BVO ECs.

#### Evaluation of genome editing

##### Fragment analysis

For quantification of fraction genome edited 409-B2 iCRISPR cells, the concentration of the two fragments generated by the T7 endonuclease and the main amplicon were measured on Bioanalyzer. The fraction of edited cells was calculated with the following formula:

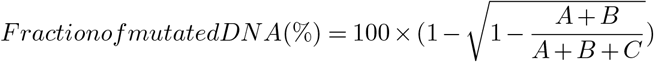

where A and B are the concentrations of the T7-generated fragments, and C is the concentration of the main amplicon.

#### Signaling pathways modulation of blood vessel organoids

##### Quantification of RT-qPCR results from pharmacologically perturbed blood vessel organoids

After initial RT-qPCR of all samples (all reactions in duplicates or quadruplicates), an additional biological replicate of each of the top five candidate conditions and control was cultured, and subsequently analyzed via RT-qPCR as described above (all reactions in duplicates). Unsuccessful RT-qPCR reactions and reactions with large a difference between the technical replicates were excluded from the final analysis (see Table S7 for Cq values, other metrics output from the qPCR and calculated. Δ ΔCt expression values for all experiments). Relative fold changes in gene expression were obtained according to the 2^-Δ ΔCt^ method (Livak and Schmittgen, 2001). In brief, first, mean quantification cycles (Cq) for each biological replicate were calculated. ΔCt and ΔCt were calculated from the mean Cq values as follows:

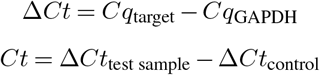

Fold differences in the gene expression of our test samples in relation to the control (original protocol) samples are then represented as 2^-Δ ΔCt^.

## Supplementary tables

**Table S1. Sample information and scRNA-seq and scMultiome data quality check. Related to all main figures**. The table provides an overview of all scRNA-seq and scMultiome experiments in this study, including sample name, number of cell pre- and post-quality control steps, mean reads per cell, median UMIs per cell, low and high cutoff for number of detected genes and high cutoff for the percentage of mitochondria transcripts.

**Table S2. Highly variable and differentially expressed genes in blood vessel organoids (BVOs). Related to figures 1 to 5** This table provides a list of 1) differentially expressed genes (DEGs) for each BVO cluster across their developmental time course; 2) highly variable genes in all scRNA-seq transcriptomic data generated in this study; 3) details about BVO endothelial cell differentiation pseudotime analysis.

**Table S3. Gene and pathway name abbreviations. Related to all main figures**. This is a list of gene and pathway name abbreviations and their full names.

**Table S4. Marker peaks of scMultiome data from day 7 BVOs. Related to Fig. 2.** This table contains the marker peaks across all peak clusters from the scATAC-seq component of the scMultiome data of BVOs at day 7.

**Table S5. Gene regulatory network (GRN) inference in BVOs. Related to figures 2, 5 and 6**. This table provides details of 1) the GRNs governing cell type identity in BVOs at day 7 (reconstructed from scMultiome data using Pando); 2) the GRN controlling the transcriptomic changes during BVO development and upon BVO transplantation (reconstructed from single-cell transcriptomic data using SCENIC); 3) signaling pathway scores in single-cell transcriptomic data from pharmacologically perturbed BVOs.

**Table S6. Experimental details and differential gene expression analysis of pooled genomic perturbation in BVOs. Related to Fig. 3.** This table provides a 1) list of transcription factors (TFs) and receptors which have been used in the CROP-seq experiments; 2) sequences of all gRNA (three per gene) used, including 8 non-targeting (dummy) gRNAs; 3) PCR primers used for gRNA validation using the T7 Endonuclease I assay; 4) This table provides differentially expressed genes (DEGs) between mural and endothelial cells with different detected gRNAs.

**Table S7. Experimental details about pharmacological perturbations in BVOs. Related to Fig. 5**. This table contains 1) details about BVO media recipes and pharmacological perturbation of all conditions and control samples; 2) sequences of qPCR primers used for assessing the effect of signaling modulation on BVOs; 3) results of RT-qPCR of the 5 chosen signaling perturbation conditions and control BVOs.

**Table S8. Vascular anomalies and diseases. Related to Fig. 6.** A list of vascular anomalies and diseases, including phenotype MIM number, details of the type of mutation, original publication and review publication references.

Table S9. qPCR primers for enrichment of scar and barcode regions of cDNA from scRNA-seq experiments. Related to Fig. 1.

## Supplementary videos

**Video 1. Time lapse movie of confocal stacks through developing blood vessel organoids (BVOs). Related to Fig. 1.** BVOs at day 3, 4, 5, 6, 7, 14, 21 stained for CD31 (endothelial cells, magenta), PDGFR-β (mural cells, yellow) and Collagen IV (basement membrane, cyan).

**Video 2. Time lapse movie of fluorescent sprouting blood vessel networks derived from induced pluripotent stem cells expressing a ubiquitous membrane-RFP-tag. Related to Fig. 1.** The movie starts at 108 h (4.5 days) after embedding in an extracellular matrix and finishes 78 h later.

**Video 3. Time lapse brightfield movies of pharmacologically perturbed sprouting blood vessel networks. Related to Fig. 5.** Scale bar: 200 µm.

## Supplementary figures

**Figure S1.**
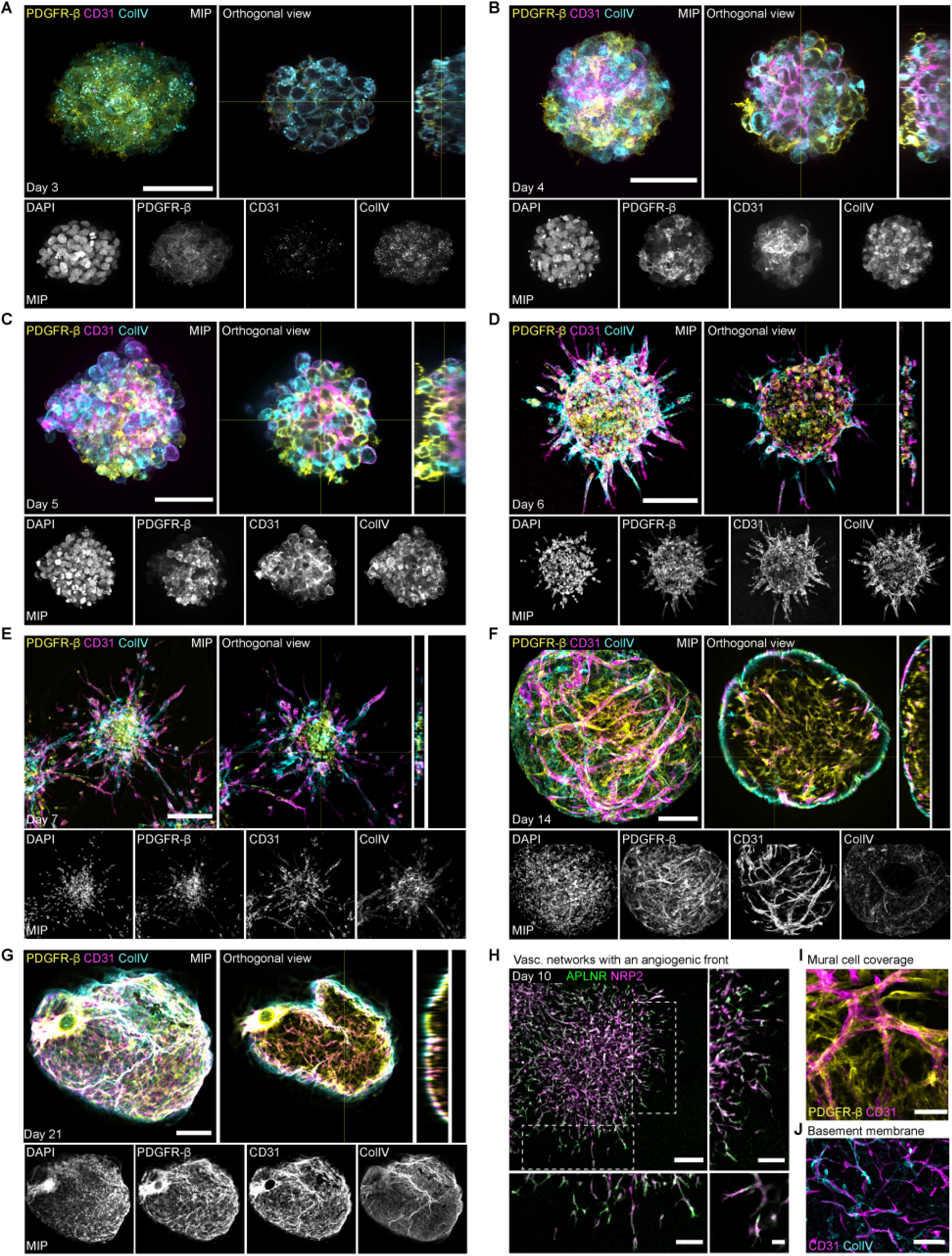
Immunohistological characterisation of hBVOs throughout development. Related to Fig. 1. (A-G) Immunostaining of whole-mount hBVOs between day 3-21 of differentiation detecting PDGFR-β (yellow, mural cells), CD31 (magenta, endothelial cells) and Collagen IV (ColIV, cyan, basement membrane). Image in the top left of each figure group is a maximum intensity projection (MIP), followed by an orthogonal view (middle) and an x-axis view (right). The bottom row of each figure group presents single channel images of the three markers and DAPI (nuclei). Scale bars: 50 µm (A-C), 100 µm (D), 150 µm (E, F), 200 µm (G). (H) Endothelial cells (NRP2, magenta) with inserts showing tip cells enriched in APLNR (green). Scale bars: 200 µm (top left image), 100 µm (top right and bottom left inserts), 25 µm (bottom right). (I) Immunostaining of day 14 whole-mount hBVO demonstrating tight pericyte coverage (PDGFR-β, yellow) of endothelial cells (CD31, magenta). Scale bar: 50 µm. (J) Endothelial cell networks (CD31, magenta) embedded in a basement membrane (Collagen IV, cyan). Scale bar: 150 µm.

**Figure S2.**
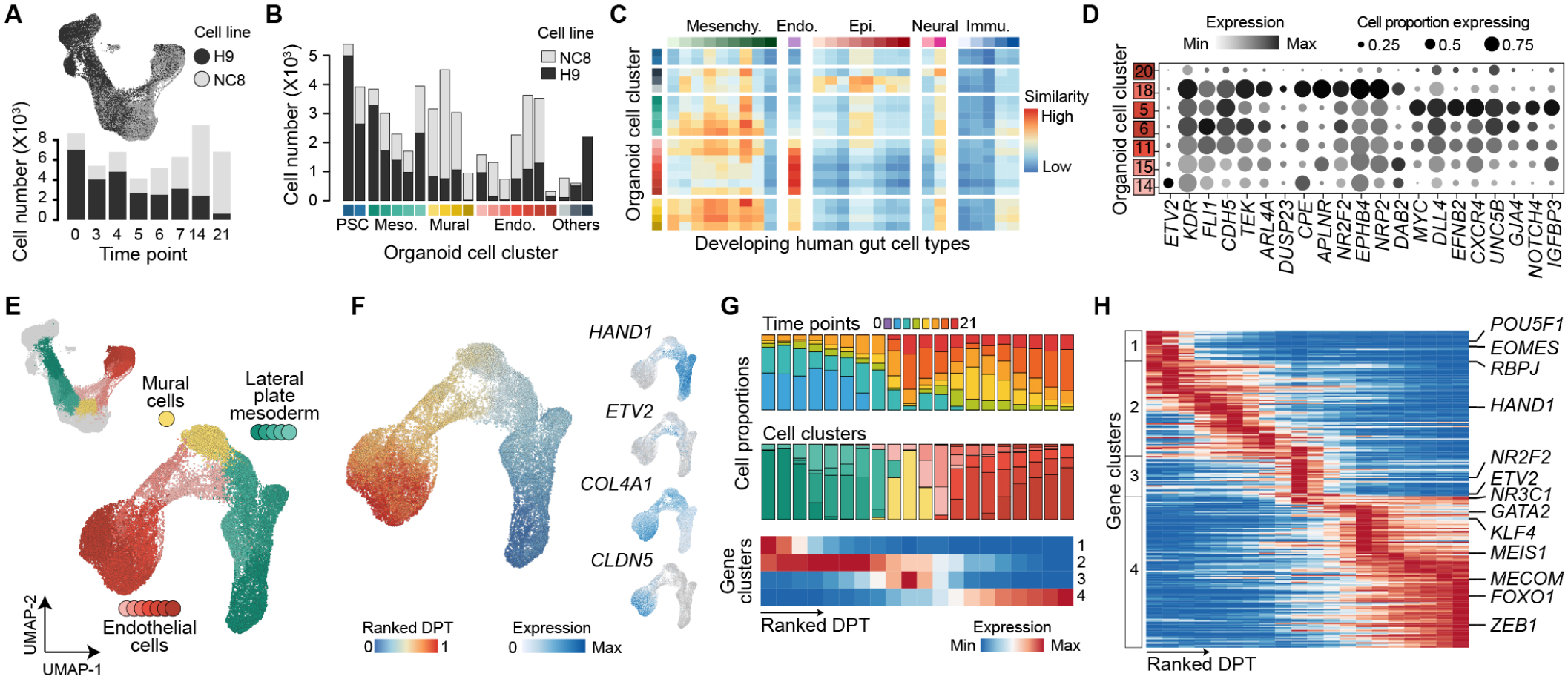
Characterization of developing hBVOs transcriptomic atlas. Related to Fig. 1. (A) UMAP embedding of single-cell transcriptomic time course colored by cell line - the embryonic stem cell line H9 (black) and the induced pluripotent stem cell line NC8 (gray). Number of cells (y-axis) per time point (x-axis) per cell line (color) used for single-cell transcriptome analysis. (B) Number of cells (y-axis) per cell cluster (x-axis colors) colored by cell line. (C) Heatmap representing the similarity of hBVO cells throughout differentiation to cells of the human developing gut (Yu et al., 2021). (D) Dotplot showing expression (color) and proportion of cells expressing (dot size) endothelial marker genes, and genes associated with arterial or venous endothelial cell fate across endothelial cells, separated by cluster. Cells in cluster 18 are enriched in venous markers, while cells in cluster 5 have a high expression of arterial markers. (E) UMAP embedding of cells forming EC differentiation colored by cell type cluster. (F) EC differentiation pseudotime represented as UMAP embedding with cells colored by ranked diffusion map pseudotime (DPT) representing. Feature plots show expression of mesoderm (HAND1) and EC (ETV2, COL4A1, CLDN5) markers along the pseudotime trajectory. (G) Proportion of cells from each time point (top) and cell type cluster (middle) per pseudotemporal bin and hierarchical clustering of pseudotemporal bins (bottom). The pseudotime trajectory was evenly split into 20 bins, every bin containing the same number of cells. (H) Hierarchical clustering showing gene markers of all pseudotemporal bins.

**Figure S3.**
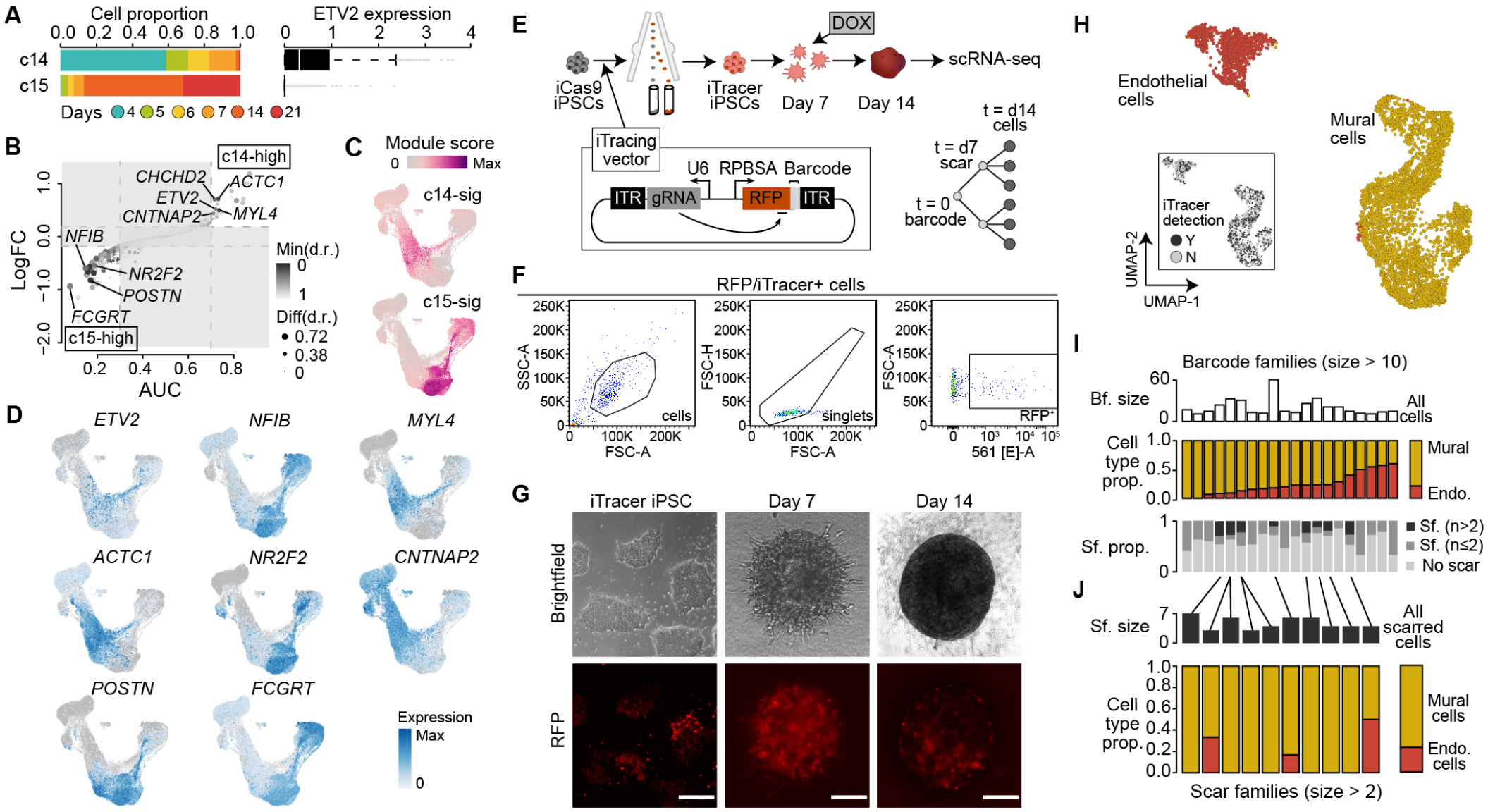
Lineage recording highlights the potential of hBVO mural progenitor cells to differentiate into endothelial and mural lineages. Related to Fig. 1. (A) Stacked bar plot showing the proportion of cells in cluster 14 and 15 (c14, c15) colored by time point and bar plot showing normalized ETV2 gene expression in the respective cluster. (B) Differentially expressed genes between c14 and c15 represented as log fold change (logFC) in gene expression versus area under the receiver operator curve (AUC). (C) Feature plots showing strength of module scores for c14 and c15 signature. (D) Feature plots showing expression DEGs between early (c14) and late (c15) EC progenitors. (E) Schematic of the iTracer experiment. A doxycycline inducible Cas9 line (iCas9) was transfected with the iTracer construct containing RFP. RFP+ cells were sorted and expanded prior to differentiation into hBVOs. Organoids were treated with a 24-hour doxycycline (DOX) pulse at day 7 to induce Cas9 and generate scars within the C-terminus of the fluorescent protein. At day 14 of differentiation, hBVOs were dissociated and scRNA-seq was performed. Regions of the scars and barcodes were further amplified and sequenced in addition. (F) FACS plots of sorting scheme used to isolate iTracer positive cells. Cells were gated first based on the main population to avoid debris, followed by gating for single cells excluding doublets, and lastly sorted for RFP positive cells. (G) Brightfield and fluorescence images of iTracer iPSCs and BVOs generated from them. Scale bars: 500 µm. (H) UMAP embedding of scRNA-seq data from day 14 hBVOs with cells colored by cell type. Insert in bottom left presents cells colored by the presence (Y) or absence (N) of iTracer detection. (I) Top: Bar plot of number of cells per barcode family for all barcode families containing more than 10 cells. Middle: Proportional stacked bar graph of cell type (mural or endothelial cell) proportion for each barcode family presented above. Bottom: Proportional stacked bar graph of scar family presence in the barcode families. (J) Top: Bar plot of number of scars in each of the scar families containing more than 2 scars. Bottom: Proportional stacked bar graph of cell type (mural or endothelial cell) proportion for each barcode family presented above.

**Figure S4.**
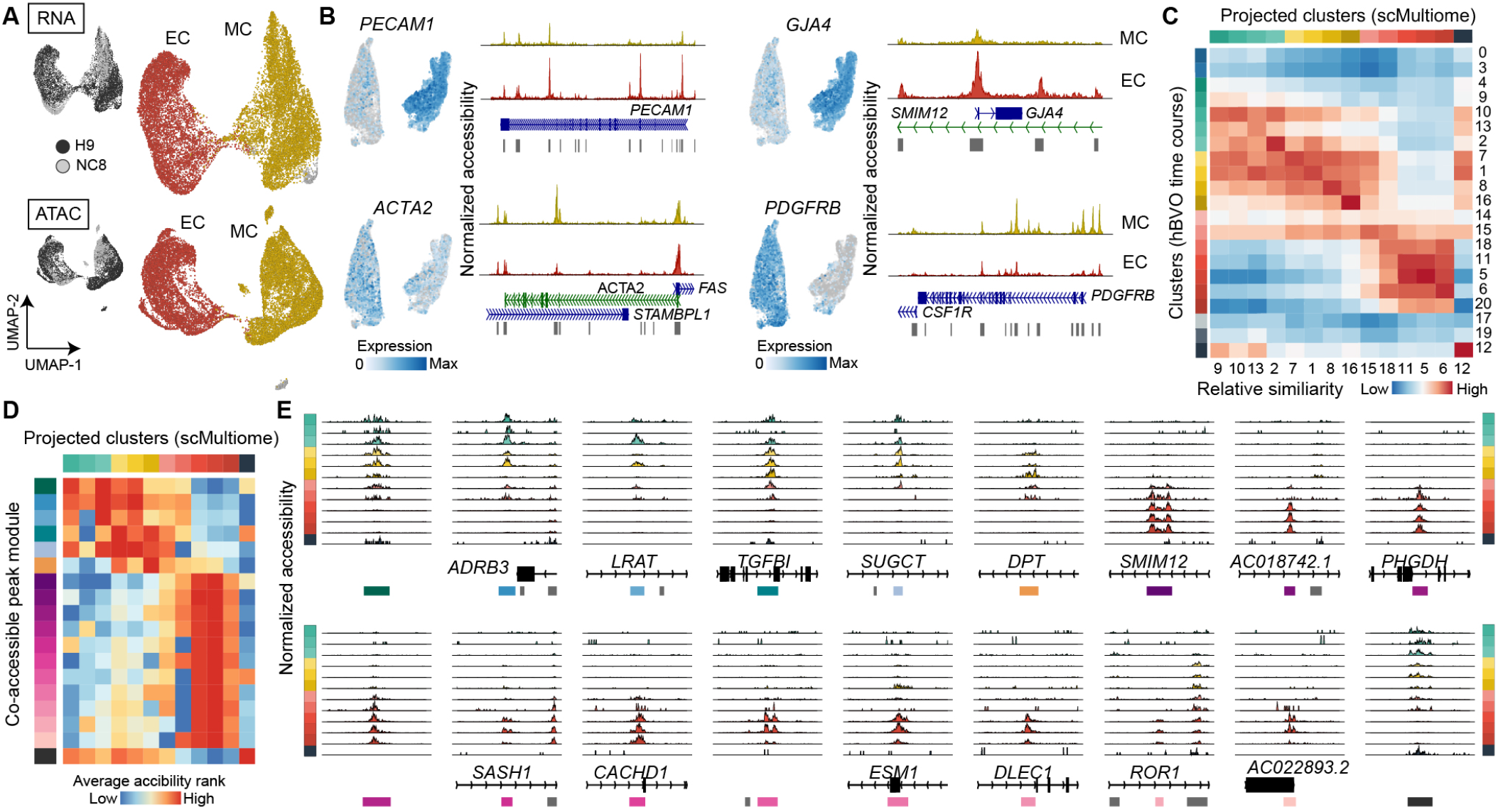
Single-cell multiomic analysis of 7-day-old BVOs reveals tissue heterogeneity. Related to Fig. 2. (A) UMAP embedding of integrated scRNA-seq or scATAC-seq of day 7 hBVOs from H9 and NC8 cells, colored by cell line (top left inserts) or by major cell type. EC, endothelial cells; MC, mural cells. (B) Feature plots and coverage plots of PECAM1 and GJA1, marking endothelial cells and ACTA2 and PDGFRB, marking mural cells. (C) Heatmap showing relative similarity between hBVO time course clusters and projected cell type clusters from the scMultiome data, based on gene expression. (D) Heatmap showing average accessibility in projected cell type clusters across co-accessible peak modules. (E) Coverage plots of example genes from each co-accessible peak module across projected cell type clusters.

**Figure S5.**
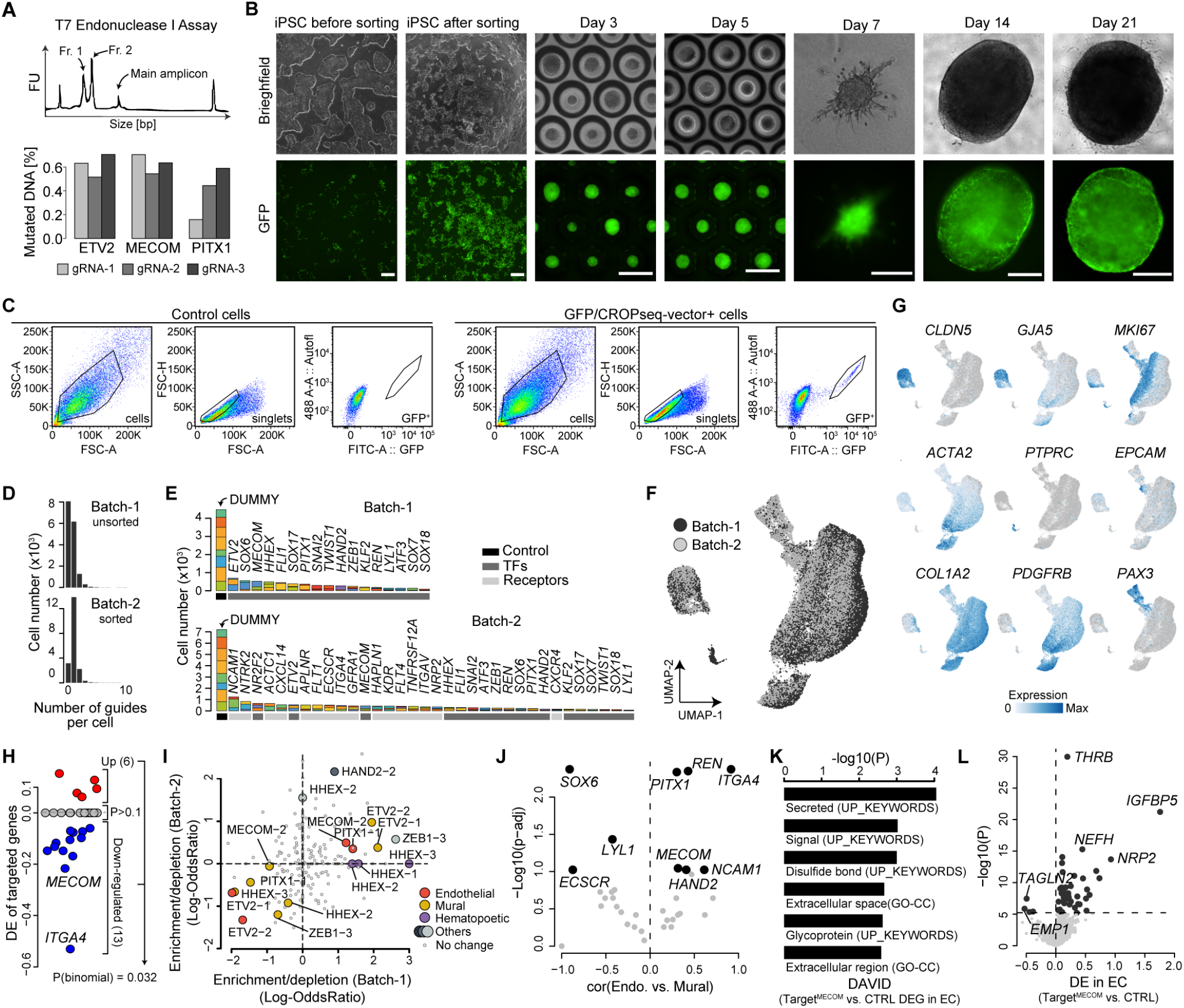
Experimental details and knock-out effect of the pooled single-cell *in organoid* perturbation experiment. Related to Fig. 3. (A) T7 Endonuclease I assay was performed with 9 gRNAs (3 different genes) to verify and quantify on-target CRISPR/Cas9 editing events in 409-B2 iCRISPR cells. After transfection with gRNA and Cas9 induction, genomic DNA from cells was amplified and heteroduplex mismatches were digested with T7 endonuclease. Bioanalyzer traces (top) of the product represent the presence of two fragments of predicted size shorter than the main amplicon (Fragment 1 and 2), indicating the successful introduction of an on-target mutation. Proportion of mutated DNA (bottom) as detected based on the T7 Endonuclease I assay. FU, fluorescence units; bp, base pair. (B) Brightfield and fluorescence images of 409-B2 iCRISPR after transduction with the CROP-seq GFP vector and during hBVO differentiation. Scale bars: 200 µm for day 7, 500 µm for all other time points. (C) FACS plots of sorting scheme used to isolate CROP-seq-vector positive cells. Cells were gated first based on the main population to avoid debris, followed by gating for single cells excluding doublets, and lastly sorted for GFP positive cells as compared to non-fluorescent controls. (D) Number of gRNAs detected per cell in batch 1 and batch 2. (E) Number of total guides targeting each gene and dummy guides detected in batch 1 and batch 2 of CROP-seq organoids. The 8 dummy guides and 3 targeting guides per gene are colored differently and the color on the x-axis shows if the guides are control ones (non-targeting), targeting transcription factors (TFs) or receptors. (F) UMAP embedding with cells colored based on the organoid batch. (G) Feature plots of marker gene expression. (H) 13 of the targeted genes were downregulated in cells where the corresponding gRNA was detected, while 6 were upregulated and the rest remained unchanged. (I) Enrichment/depletion of guides in different cell types in batch 1 versus batch 2. (J) Correlation between the knock-out effect (on gene expression) of particular genes in endothelial versus mural cells. (K) Examples of functional enrichment for DEGs in ECs with detected MECOM gRNAs with DAVID. (L) Differentially expressed genes (DEGs) in ECs in which gRNAs targeting MECOM were detected versus cells with dummy gRNAs detected.

**Figure S6.**
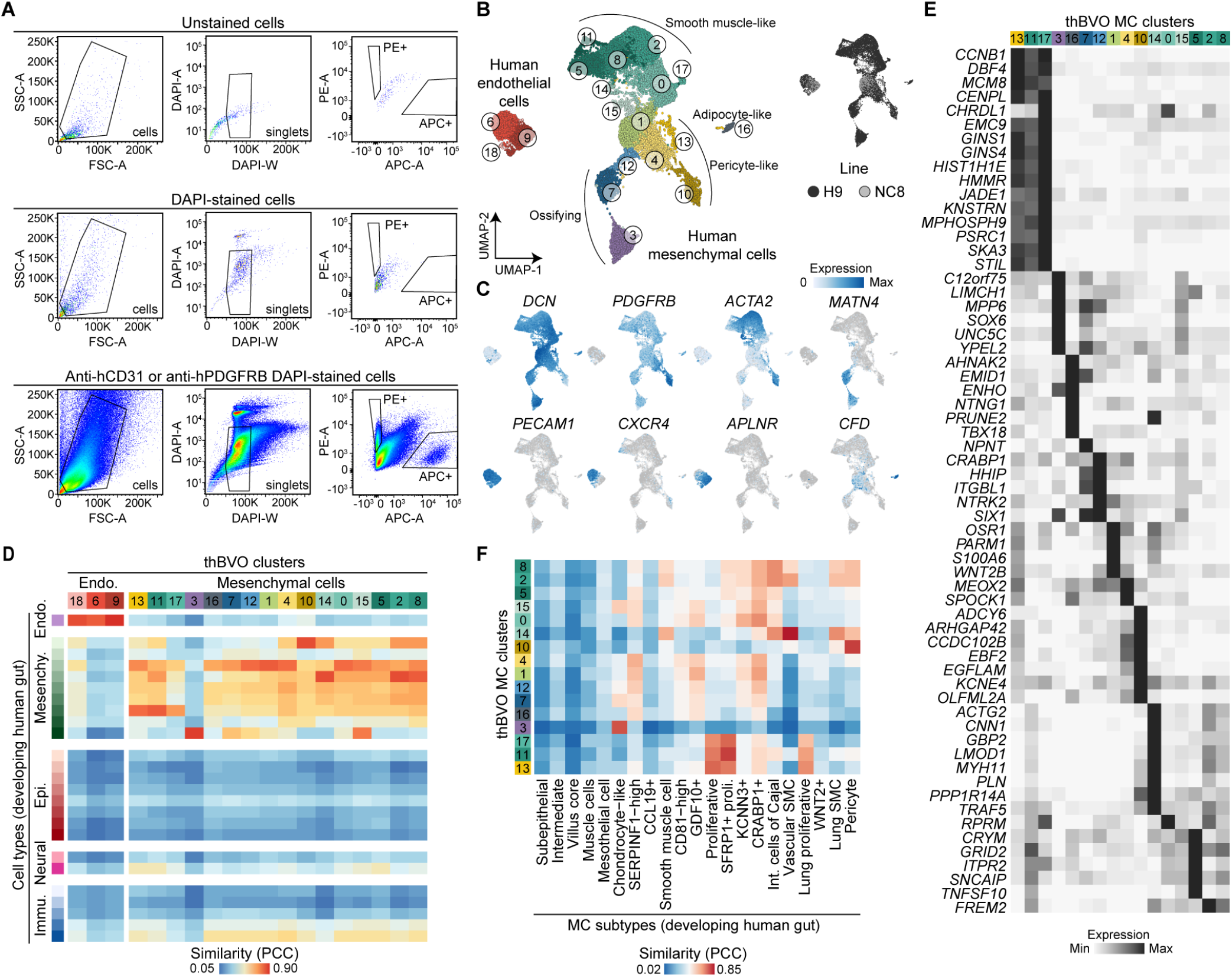
Transplantation of human BVOs into immunocompromised mice can lead to expansion of off-target fates. Related to Fig. 4. (A) FACS plots of sorting scheme used to isolate APC-hCD31 and PE-hPDGFR-positive cells. Cells were gated first based on the main population for single cells excluding doublets (plot at the top), followed by gating based on DAPI staining to avoid debris and dead cells (plot in the middle), and lastly sorted for APC (human ECs) and PE (human pericytes) positive cells as compared to non-fluorescent controls (plot at the bottom). (B) UMAP embedding of transplanted hBVO (thBVO) CD31^+^ and PDGFR-β^+^ with cells colored by cluster, or by cell line (ESCs, H9 and iPSCs, NC8). (C) Feature plots showing expression of endothelial (CLDN5, CXCR4, APLNR) and mesenchymal (DCN, PDGFRB, ACTA2, SIX1) marker genes in thBVOs. (D) Heatmap representing the similarity of thBVO cells to cells of the human developing gut (Yu et al., 2021). PCC, Pearson correlation coefficient. (E) Heatmap showing averaged expression of cluster markers genes in mesenchymal cells (MC) from thBVOs. (F) Heatmap showing the similarity of MC thBVO cell clusters to MC subclusters of the human developing gut.

**Figure S7.**
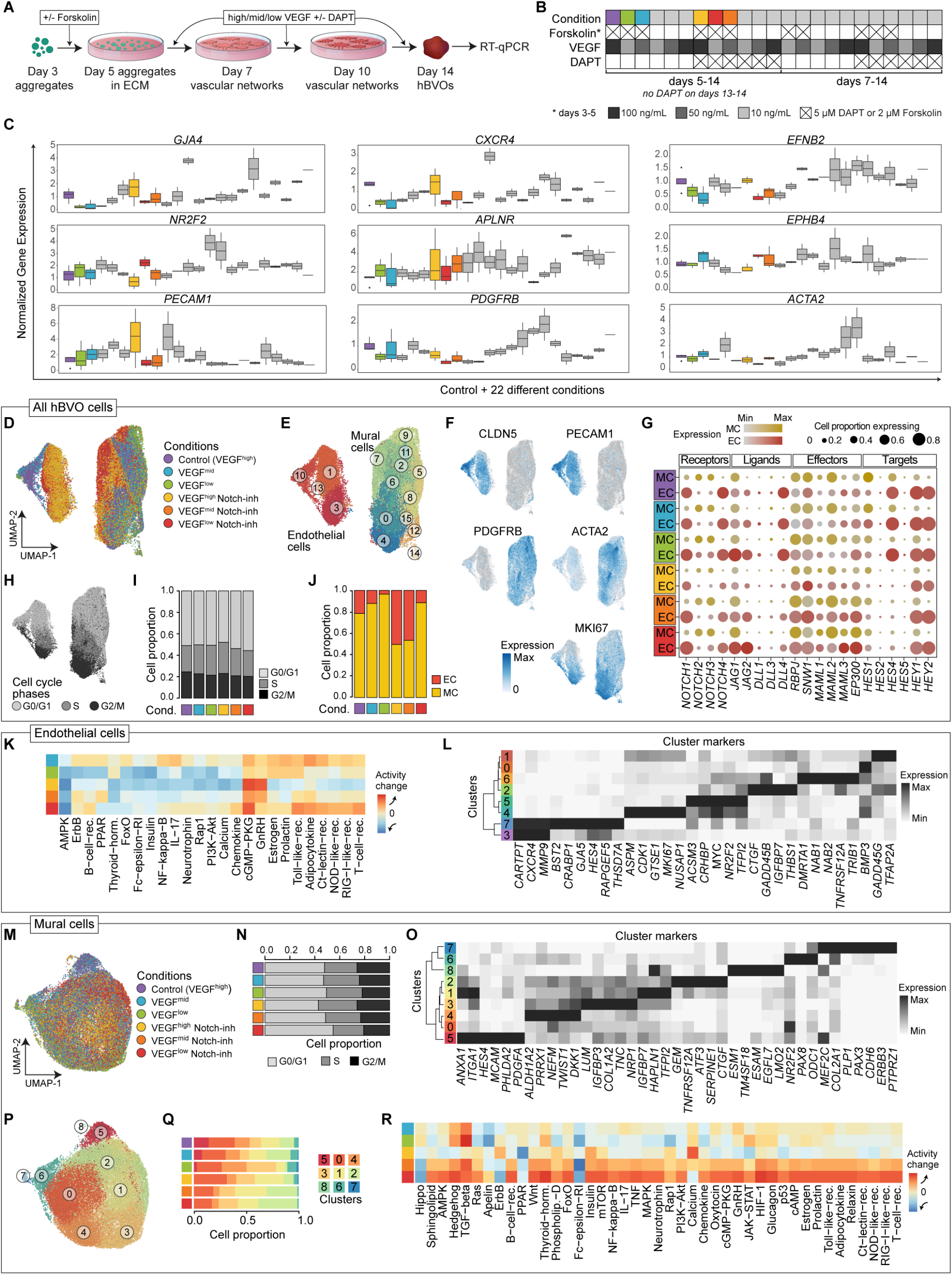
Experimental design of the pharmacological screen in BVOs and effects on EC and mural cell differentiation. Related to Fig. 5. (A) Schematic of signaling perturbation experiments. (B) Overview of signaling perturbations. (C) RT-qPCR of endothelial subtype and mural cell markers showing gene expression normalized to GAPDH in day 14 hBVOs. Here 22 different conditions were tested (see Table S7) and only the conditions for which follow up experiments were performed are in color. Data are represented as log fold change, showing distribution and median. n = 2 to 3. (D, E) UMAP embedding of hBVOs colored by signaling pathway perturbation (D) or by cell type cluster (E). (F) Dot plot showing average gene expression of Notch receptors, ligands, effector and target genes in endothelial (EC) and mural (MC) cells across conditions. Dot size indicates the percentage of cells expressing the gene. (G) Feature plots showing gene expression of endothelial (CLDN5, PECAM1), mural (PDGFRB, ACTA2) and proliferating (MKI67) cell marker genes. (H) UMAP embedding of hBVOs colored by cell cycle phase. (I) Proportion of hBVO cells per cluster colored by cell cycle phase (top) or by major cell type (bottom). (J) Immunohistochemical staining detecting TIE1 (magenta) and PDGFR-β (green) in cells across control and pharmacologically perturbed hBVOs at day 10 of differentiation. Scale bar: 250 µm. (K) Heatmap of signaling pathway activity across signaling pathway perturbation normalized to control cells in ECs only. (L) Heatmap showing average gene expression of cluster marker genes in ECs. (M) UMAP embedding of MCs colored by signaling perturbation condition. (N) Proportion of mural cells per cluster colored by cell cycle phase. (O) Heatmap showing average gene expression of cluster marker genes in MCs. (P) UMAP embedding of MCs colored by cell type cluster. (Q) Proportion of mural cells per cluster colored by perturbation condition. (R) Heatmap of signaling pathway activity across signaling pathway perturbation normalized to control cells in MCs only.

